# Male Guinea baboons are oblivious to their females’ whereabouts

**DOI:** 10.1101/2022.07.20.500821

**Authors:** Dominique Treschnak, Dietmar Zinner, Julia Fischer

**Affiliations:** Cognitive Ethology Laboratory, German Primate Center, Göttingen, Germany; Department of Primate Cognition, Georg-August-University of Göttingen, Göttingen, Germany; Leibniz ScienceCampus Primate Cognition, Göttingen, Germany

## Abstract

In group-living species, evolution puts a premium on the ability of individuals to track the state, whereabouts, and interactions of others. The value of social information might vary with the degree of competition within and between groups, however. We investigated male monitoring of female location in wild Guinea baboons (*Papio papio*). Guinea baboons live in socially tolerant multi-level societies with one-male-units comprising 1-6 females and young at the core. Using field playback experiments, we first tested whether male Guinea baboons (N=14) responded more strongly to playbacks of associated vs. non-associated females, which was the case. In the second and core experiment, we tested whether males (N=22 males, N=62 trials) keep track of the whereabouts of associated females by playing back unit females’ calls from locations that were either consistent or inconsistent with the actual position of the female. Contrary to predictions, males responded equally strongly in both conditions. While males seem to recognize their females by voice, they might lack the attention or motivation to track their females’ movement patterns. These results reinforce the view that the value of social information may vary substantially with the distribution of power in a society. While highly competitive regimes necessitate high attention to deviations from expected patterns, egalitarian societies allow for a certain degree of obliviousness.

## Introduction

Knowledge about conspecifics and their relationships guides social decision-making in many group-living animals. The use of such social knowledge is documented for a large number of species, ranging from simple and more complex forms of individual recognition (Wiley, 2013) to the assessment and monitoring of stable or transient social attributes of group members, like kinship, rank, or bond strengths. Such knowledge extends not only to an individual’s direct associations but also to third-party relationships (Seyfarth & Cheney, 2015). When navigating the social environment, knowledge about previous interactions with group members, the capabilities of potential partners or competitors, and the nature and quality of relationships between others, aids in predicting the outcomes of future interactions and allows to act strategically. For example, spotted hyenas (*Crocuta crocuta*) joining into dyadic fights mainly support the dominant individual and are subsequently also more likely to attack relatives of the subordinate (Engh et al., 2005). Pinyon jays (*Gymnorhinus cyanocephalus*) assess their relative rank difference to strangers by observing them in encounters with known individuals (Paz-y-Miño C et al., 2004). Tonkean macaques (*Macaca tonkeana*) respond more strongly to conflicts between strongly bonded individuals (‘friends’) compared to non-friends (Whitehouse & Meunier, 2020).

Besides kin and allies, mating partners are of particular value to an individual. Males compete not only for access to females (Clutton-Brock & Parker, 1992; Clutton-Brock & Vincent, 1991); they are also under selection to monitor the state and behaviour of females. Males may increase their reproductive success by assessing suited mating partners (Davies et al., 2020) or mating opportunities (Balsby & Dabelsteen, 2005; Crockford et al., 2007). In many species, females become the centre of male attention when they approach the fertile phase of their reproductive cycle. In contrast, in species where males and females form long-lasting bonds as in monogamous (Birkhead & Møller, 1995) or polygynandrous species (e.g., plains zebras (*Equus burchellii*) (Rubenstein & Hack, 2004), hamadryas baboons (*Papio hamadryas*) (Swedell & Plummer, 2012)), males are permanently incentivised to monitor and control associated females’ whereabouts and interactions with other group members.

We tested male knowledge of female whereabouts in wild Guinea baboons (*Papio papio*). The species lives in multi-level societies. At the core are one-male units consisting of one primary male, one to six associated females, and their offspring. Bachelor males may be associated with several such units (Dal Pesco et al., 2021). Several units form a party, which in turn aggregate into gangs (Patzelt et al., 2014). Females associate with one primary male and show mate fidelity (Goffe et al., 2016). Still, in contrast to hamadryas baboons, they also enjoy spatial freedom, i.e., they may spend considerable time away from their male (Goffe et al., 2016). Females may transfer to other males between all levels of the baboon society. Transfers have even been observed for females while pregnant or with a dependent infant. We hypothesized that primary males keep track of the movement patterns of their associated females, as proximity or interactions between associated females and other males could indicate potential transfer intentions of their females to primary males. To test this hypothesis, we conducted a playback experiment (Fischer et al., 2013), in which we presented female grunts from a location that was either consistent or inconsistent with the actual position of the female. We made use of the violation-of-expectation paradigm and presented the animals with a physically impossible scenario, similar to Townsend et al. (2012). We tested a male immediately after the female had left him and assumed that he would have noticed the direction in which she disappeared. We predicted that males would show ‘signs of surprise’, meaning a stronger response, when they were confronted with information that the female was in an unexpected – indeed physically impossible – location compared to their response when the female’s vocalisation came from the direction into which she had recently disappeared. In a preparatory experiment, we tested the prerequisite that males can recognise their associated females by voice. We tested if males respond more strongly to the vocalizations of females from their unit compared to the vocalizations of females from another unit but the same party. We predicted that males would show stronger responses when presented with vocalisation from unit females.

## Methods

The experiments took place between January 2019 and August 2021 at the Centre de Recherche de Primatologie Simenti in the Niokolo-Koba National Park in Senegal, a field station maintained by the German Primate Center (see Fischer et al., 2017 for details). The study population comprised ∼ 200 individually identified Guinea baboons that belonged to three parties, with a varying number of reproductive units (between 15 and 25 per year across three parties) suitable for the experiments. The baboons are habituated to the presence of researchers and allow approaches within a few meters without signs of disturbance.

For the experimental stimuli, we recorded ‘grunt’ vocalisations of sub-adult and adult females during their non-receptive phase, i.e., the females did not show any swelling of their anogenital skin as a sign of high ovulation probability. Individual females were chosen opportunistically based on the availability of high quality recordings. Grunts are the most frequently occurring vocalisation in Guinea baboons (47.8 ± 30.1 call elements/h/individual) and are mainly produced in affiliative contexts (Maciej, Ndao, et al., 2013). To produce high-quality experimental stimuli, we selected only recordings with a high signal-to-noise ratio, i.e., a large difference between signal amplitude and the amplitude of other background sound sources and no other sounds overlaying the individual grunts. We inserted silent segments between individual grunt elements and normalised call amplitudes to a percentage of their dynamic range (65 - 90%). Female grunt vocalisations show individual differences in structural characteristics, such as the length and number of elements within a grunting bout. While we wanted to maintain these inter-individual differences, we also wanted to use stimuli that did not vary too much in their structure, potentially influencing the responses of males independently of the test condition. We, therefore, set limitations for the total length of a grunt sequence (duration from the start of the first grunt to the end of the last grunt), number of grunts per sequence, and the total grunt duration within a sequence (sum of the length of all individual grunts within a sequence). The final stimuli had an average total length of 2.23 s (2.12 –2.49 s), an average number of six grunt elements per sequence (4–7), and an average total grunt duration per sequence of 0.58 s (0.48 – 0.74 s). We measured the sound pressure level for each stimulus at a distance of 10 m (comparable to experimental conditions). We also controlled whether the stimuli sounded subjectively similar to actual vocalisations of female baboons (figure 1).

**Figure 1:**
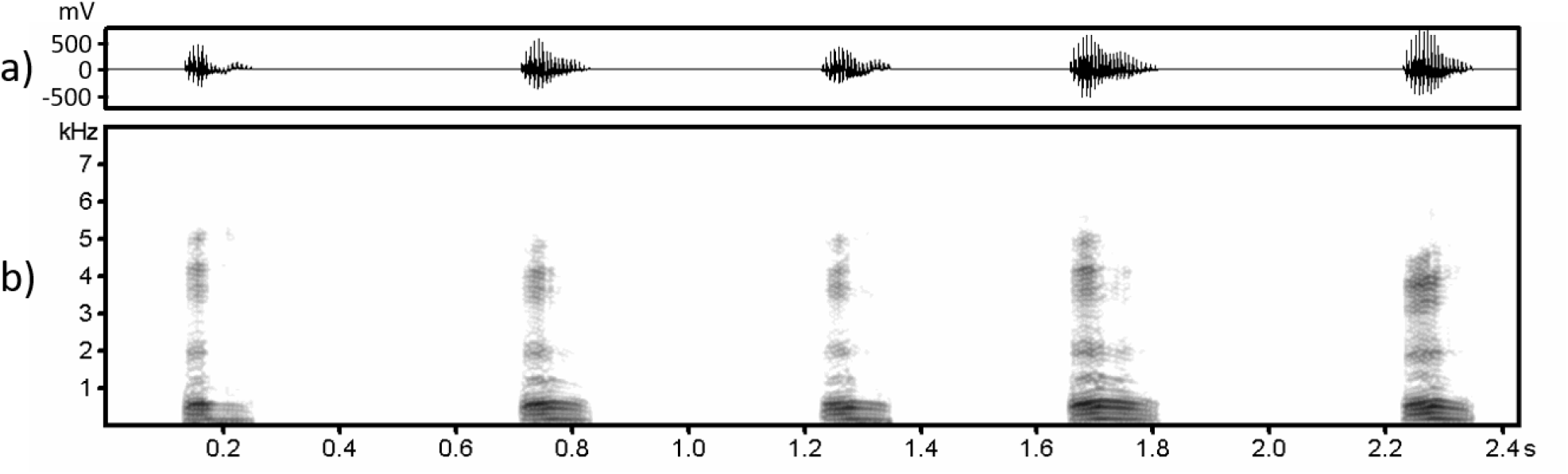
Example of experimental stimulus. a) Waveform (envelope) of the call amplitude changing over time. b) Spectrogram depicting the distribution of different amplitudes (shades of grey) over the frequency spectrum and over time. FFT length = 512, Hamming window, overlap 93.75 %, sampling frequency = 16 kHz, time resolution = 2 msec.

For the recordings, we used a solid-state recorder (Marantz PMD661 MKII, Marantz, Kanagawa, Japan) with Sennheiser directional microphones (K6 power module with ME66 recording head, Sennheiser Electronic KG, Barleben, Germany) and Rycote windshields (Rycote, Gloucestershire, UK) with a sampling frequency of 44.1 kHz and 16-bit resolution. We only used recordings taken from a distance < 5 m to the animal to avoid effects of signal attenuation (Maciej et al., 2011)). We used Avisoft-SASLab Pro 5.2 (Avisoft Bioacoustic, Glienicke, Germany) and Audacity 3.1 (Audacity Team, https://audacityteam.org) to analyse and prepare the playback files.

In Experiment 1 (individual recognition), we presented males with calls from a female from their unit (*unit-female* condition) and a female from another unit (*non-unit-female* condition). Trials were separated by at least five days and conducted only when females were non-receptive. Once the female whose call was to be played back was not visible to the subject, a loudspeaker was positioned at a 90° angle to the left or right of the male depending on the actual position of the female, and the stimulus presented. Male responses were video recorded for three minutes after the onset of the stimulus. We conducted 28 playback trials testing 14 primary males.

In Experiment 2 (spatial monitoring), we tested males in a within-subject design and presented grunts from a unit-female on two occasions separated by at least seven days. As above, trials were conducted only when females were non-receptive. In the *consistent* condition, the loudspeaker was hidden in a location matching the actual direction of the departed female. In contrast, in the *inconsistent* condition the loudspeaker was hidden in the opposite direction (i.e., the angle in direction between the positions of loudspeaker and female was ∼180°), presenting an impossible scenario (figure 2). A male was tested after he had been near a unit-female, she had then walked away and was no longer in sight. We aimed to conduct trials within 180 s of the female being out-of-sight to ensure that the female would not be able to reach the location of the loudspeaker (median time out-of-sight: 70 s, range 8 s – 273 s). A loudspeaker was then hidden in vegetation, at a 90° angle to the left or right of the male and a distance of approximately 10 m. Male responses were video recorded for 10 min. after the onset of the stimulus. Throughout the experimental trials, only the researcher who video-recorded the response of the male from a distance of 3 to 7 m was visible to the male. We only conducted experimental trials in generally calm periods, i.e., not during aggressive episodes among group members and not on days where groups encountered predators or other disturbances occurred. Males were only tested when they were resting or feeding, not being in direct contact or interacting with other adult baboons and if no other baboon was between the loudspeaker and the male. We conducted 62 playback trials with 22 primary males. Nine of these males were tested twice with the call of a different female (average time between first and second run: 43 weeks (min: 3, max: 100)).

**Figure 2:**
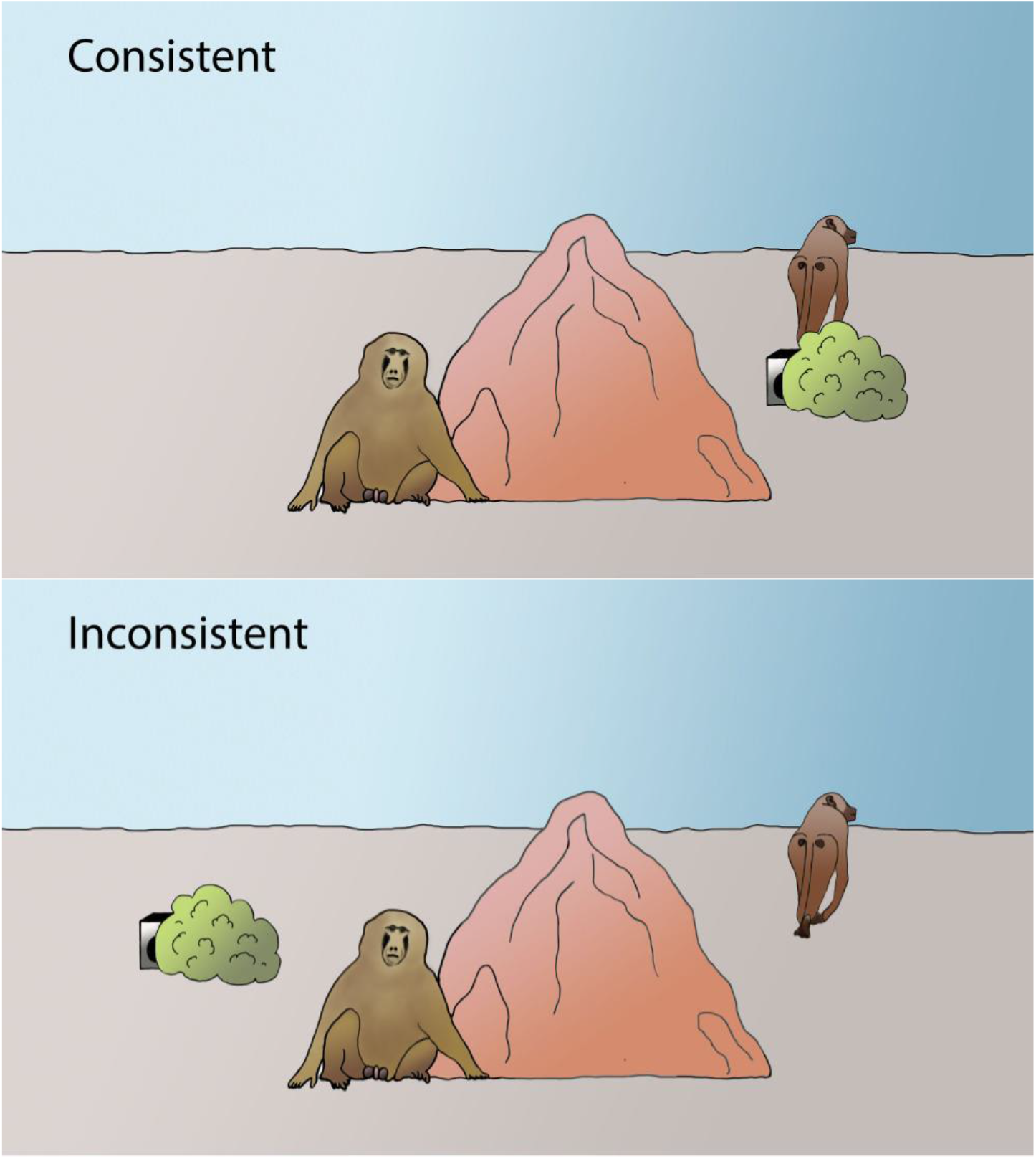
Set-up experiment 2 (spatial monitoring). In the consistent condition, a loudspeaker is positioned close to the location where the female has been before leaving, in the inconsistent condition, the loudspeaker is placed in the opposite direction in respect to the male’s position.

The sequence in which experimental conditions were presented was counterbalanced and then randomly assigned to individuals. Males that were tested with a second female in experiment 2 were exposed to the experimental conditions in the opposite order in their second run. We could not control the direction from which stimuli were presented to males as the positions of male, female and the trial condition dictated the experimental setup. We only used recordings from females that were non-receptive at the time of the recording to avoid a potential influence of female reproductive state. Also, we only conducted the trials when females were non-receptive.

For the playback, we first used a DAVIDactive loudspeaker with an integrated battery (VISONIK, Berlin, Germany) connected to a handheld solid-state recorder (Marantz PMD661 MKII, Marantz, Kanagawa, Japan) by cable. In 2021, we switched to a wireless loudspeaker (Sonos Move, Sonos, Santa Barbara CA, US) connected to a Gigaset GX290 smartphone (Gigaset, Bocholt, Germany) via a portable Wifi Network from a second Gigaset GX290 smartphone. With the new wireless set-up, experimental trials could be conducted more efficiently. Old and new loudspeakers were accordingly adjusted to produce qualitatively comparable stimuli. We conducted 22 trials with the first set-up and 68 with the second. Videos were recorded using a Panasonic HC-X909 video camera (Panasonic Corporation, Kadoma, Japan). Sound pressure levels were, measured using a sound level meter (Voltcraft SL-400, Voltcraft, Germany).

Video recordings were coded using Solomon coder beta (András Péter, solomon.andraspeter.com) on a frame-by-frame basis (25 frames/s). We examined male responses by coding changes in their head orientation; i.e., changes between the neutral position: male faces the camera or in the opposite direction to the loudspeaker, and subsequent looks exceeding an angle of 45° towards the direction of the loudspeaker or away from it. We measured the duration of the first-look and the latency to respond. We measured the onset of the first responses for all trials and examined the histogram of latencies blind to the experimental condition searching for a latency which separates putative actual responses with a short latency from later responses which might have happened for reasons other than the playback. We settled on a cut-off criterion for responses to be counted as valid if they occurred within the first 2.5 s of presenting the stimulus (figure 3). Responses that occurred after the cut-off criterion of 2.5 s were counted as non-response, with a duration of 0 s and censored latency. As the first look in the inconsistent condition could be truncated because the male may turn his attention to look into the direction where the female was last seen, we additionally measured the total time vigilant (all looks toward the loudspeaker or actual position of the female) within 30 s after stimulus onset in the social monitoring experiment, (appendix, figure A1 (classification of responses)).

**Figure 3:**
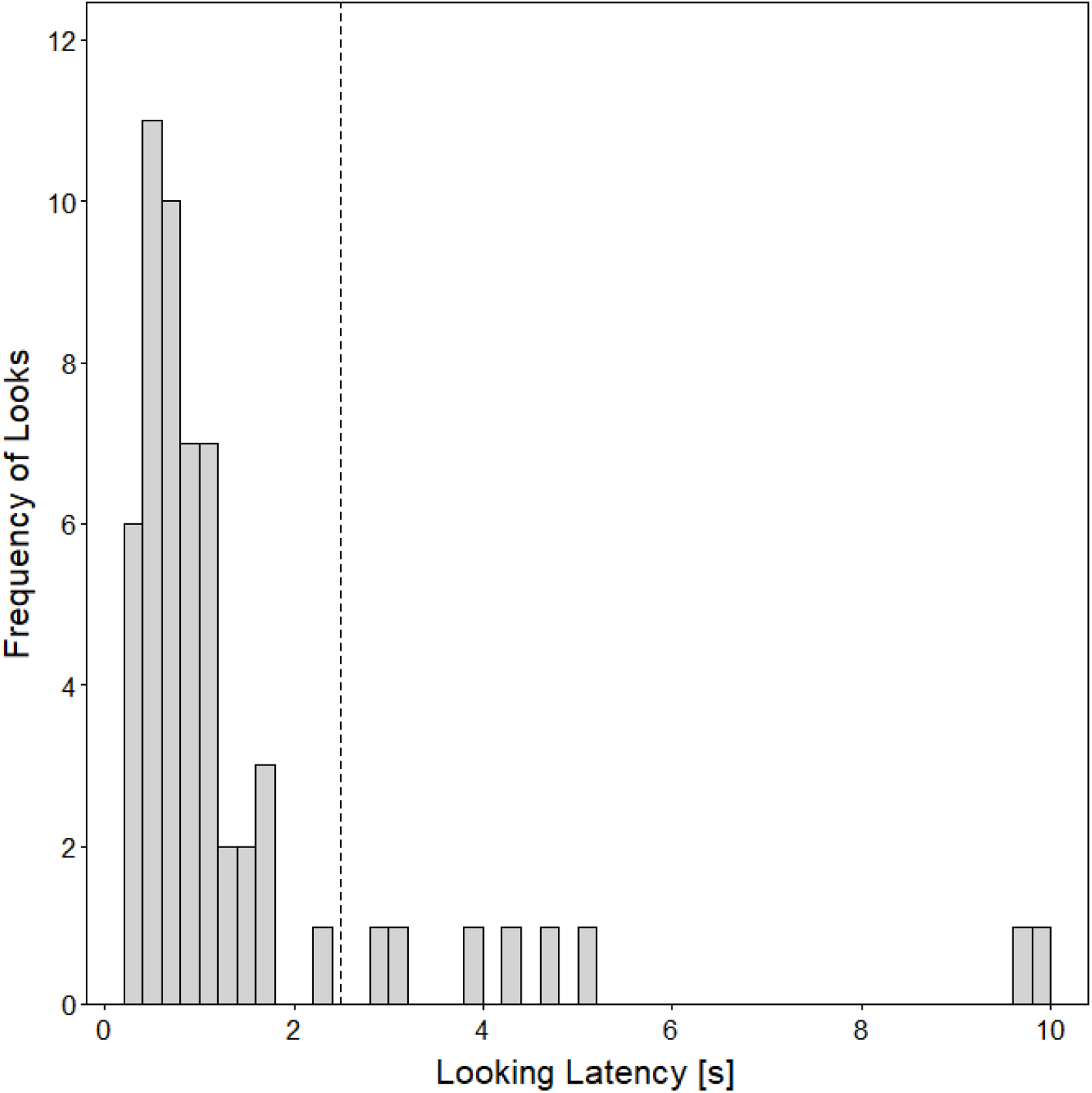
Histogram of the latency to the first response after presentation of the experimental stimulus. In 49 out of 62 trials a response occurred within the first 2.5 s. The vertical dashed line shows the selected cut-off point for valid responses at 2.5 s.

For twenty randomly selected trials (representing 22% of all trials), the video recordings were coded by a second observer blind to the general experimental setup and research question and compared to the coding results of the first author (DT). We compared the amount of correctly coded changes in head orientation (according to the 45° rule; see Methods) and the looking durations and latencies for each trial. Both coders correctly agreed on the occurrence of the first response in 19 out of 20 trials. Consecutive head orientation changes after the first response were correctly coded in 43 out of 55 instances (78%). The few head orientation changes that were not coded by both observers correctly were either relatively short glances or head movements where it was difficult to determine whether or not the 45° rule was fulfilled. As first responses could be clearly and reliably detected in almost all trials by both observers, our main criterion for reliability was the exact quantitative coding of the response. A Spearman’s rank correlation for looking duration and latency revealed high inter-observer reliability for both measures (duration: rho = 0.99, *P* < 0.001; latency: rho = 0.95, *P* < 0.001). After the reliability of the coding procedure was ascertained, DT coded and analysed the full data set.

We used a Linear Mixed model (Baayen et al., 2008) for first-look duration (experiment 1) and vigilance time, a Generalized Linear Mixed Model with gamma error structure and log link function (Baayen et al., 2008) for the duration of the first-look (experiment 2), and a survival analysis (Jahn-Eimermacher et al., 2011) for latencies. In addition to the main predictor ‘experimental condition’, we included total unit size for each male as a fixed effect to control for the influence of the number of unit-females and male identity as random intercept. To investigate the effect of the main predictor we compared full models to null models lacking the main predictor of interest (experimental condition) in the fixed effect part, using a likelihood ratio test (Dobson & Barnett, 2008). Confidence intervals of estimates and fitted values were determined using a parametric (LMM & GLMM) and non-parametric (survival analysis) bootstrap (*N*=1000 bootstraps). Responses that occurred after the cut-off criterion of 2.5 s were considered censored for survival models and entered the duration models with a length of zero. Model stability was assessed by comparing model estimates for the complete data set with estimates for data sets with levels of the random effect (subject) excluded one at a time.

Experiment 1 (individual recognition): Model validity checks for the first-look duration model revealed no obvious deviations from assumptions of normality or homoscedasticity, which allowed for the use of a Linear Mixed Model. For the first-look latencies, we used a survival analysis (Cox proportional hazard model), as this method allows for the inclusion of trials without a response and has fewer distributional assumptions. Prior to the analysis, we log-transformed (base e) first-look durations to achieve an approximately symmetrical distribution. The assessment revealed acceptable stability for the survival model (see table A2). We found instability in the range for the estimated effect of the main predictor ‘experimental condition’ in the Linear Mixed Model for first-look duration (see table A1 – A2), which could be traced back to the influence of one single trial, that, when being removed, led to a dysfunctional model with essentially zero residual variance and hence unrealistic model estimates. The observed instability can therefore be seen as an artefact and not as the effect of removing an influential case. As the same call (stimulus) could be used for different males in Experiment 1, and some calls were from the same female, we included female ID and stimulus ID as additional random intercept effects. No random slopes were included, as none were theoretically identifiable.

Experiment 2 (spatial monitoring): We used Generalised Mixed Models for the first-look duration as model validity checks revealed deviations from assumptions of homoscedasticity. We found no deviations for vigilance durations and used a Linear Mixed Model. For first-look latencies, we again calculated a survival analysis (Cox proportional hazard model). In the GLMM, ‘first-look’ durations of 0 were changed to 0.01 s to allow the use of the gamma error distribution. Vigilance durations were squared to achieve a more symmetrical distribution. The assessment revealed acceptable stability (see table A3 – A5). The GLMM showed slight underdispersion (dispersion parameter = 0.93; dispersion parameter <1 reveals underdispersion). Underdispersion can lead to over-conservative model estimates, which would, in this case, unlikely change the general interpretation of the model results. We did not calculate confidence intervals for the estimated effects in the survival model, as to our knowledge, there is no methodological approach available that allows calculating confidence intervals for Cox proportional hazard models with more than one random effect included.

After conducting the data analyses for the spatial monitoring experiment, we discussed the potential influence on male attention of the presence of other baboon parties when conducting the trials. We, therefore, formulated new models for all three outcome variables including a binary indicator indicating whether other parties were overlapping in space with the party of the tested male (other parties in proximity (mingled) = Y; no parties in proximity = N) (model estimates for all three alternative models, table A6). Information about the presence of other parties was recorded directly after the conduction of successful trials.

Analyses were carried out in R (version 4.1.1; R Core Team, 2021). GLMMs and LMMs were fitted using the function *glmer* of the R package *lme4* (version 1.1-27.1; Bates, 2015). For the Cox proportional hazard model, we used the packages *survival* (3.2-13) and *coxme* (2.2-16). Model stability and overdispersion were assessed using a function provided by Roger Mundry (see details just below). The bootstrapped confidence intervals were obtained using the function *bootMer* of the package *lme4*. We calculated test statistics and *P*-values for LMMS using the *lmer* function of the *lmerTest* package (3.1-3).

This research adhered to the ASAB/ABS Guidelines for the Use of Animals in Research (‘Guidelines for the Treatment of Animals in Behavioural Research and Teaching’, 2020). Approval and research permission were granted by the Direction des Parcs Nationaux and the Ministère de l’Environnement et de la Protection de la Nauture de la République du Sénégal (date: 22/04/2019). Research was conducted within the regulations set by Senegalese agencies as well as by the Animal Care Committee at the German Primate Center (Göttingen).

## Results

In experiment 1 (individual recognition), males responded to the playback of calls in 24 out of 28 trials. The average duration of the first response was 3.2 s ± 2.5 s (median ± IQR). Males looked longer when presented with calls from unit-females (3.4 s ± 4.6 s) compared to non-unit females (2.3 s ± 3.8 s) (figure 4a; full-null model comparison: χ^2^_1_ =8110, *P*=0.004, table A1a). The average latency of responses was 1.0 s ± 0.9 s for the unit-female and 1.3 s ±3.2 s for non-unit-females (median ± IQR). Unit size had no obvious effect on response duration or latency (Duration: *P*=0.48; Latency: *P*=0.37, table A1a, A2).

**Figure 4:**
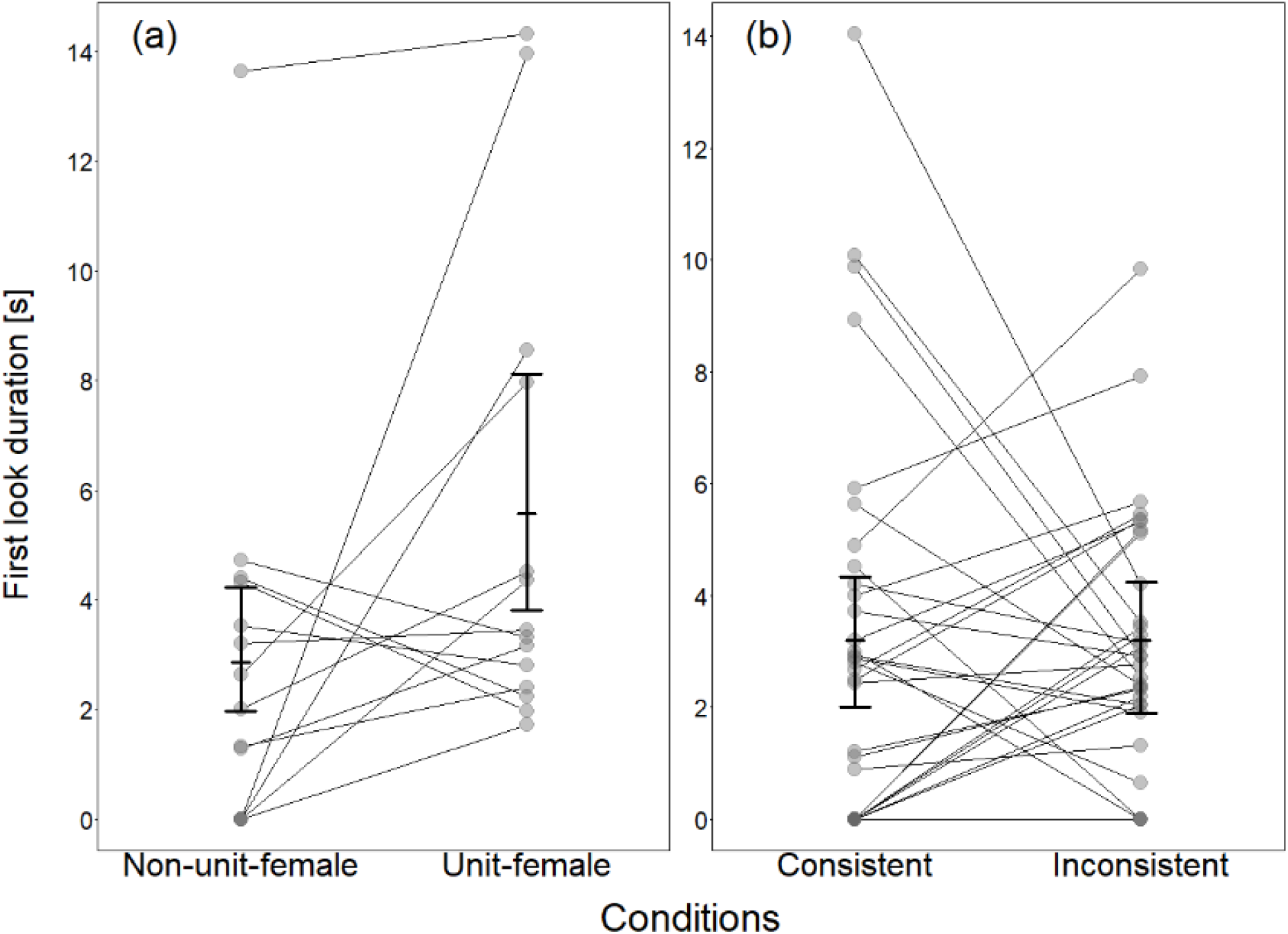
First-look duration for males in the a) individual recognition experiment and b) spatial monitoring experiment. Connected points represent data from the same individual (a: *N*=14; b: *N*=22). Thick black lines depict bootstrapped mean and 95% confidence intervals for males with average unit size.

In experiment 2 (spatial monitoring), males responded to the playback in 49 out of 62 trials (consistent condition: *N*=22, inconsistent: *N*=27). There was no obvious difference in the duration of first-looks in the consistent (2.8 s ± 4.4 s, median ± IQR) compared to the inconsistent (2.9 s ± 2.6 s) condition (figure 4b; full-null model comparison: χ^2^_1_=0.0002, *P*=0.99, table A3). There were no obvious differences in response latencies between the two conditions (consistent: 0.7 s ± 0.5 s; inconsistent: 0.8 ± 0.6 s (median ± IQR); full-null model comparison: χ^2^_1_ =1.10, *P*=0.29, table A4). There were also no obvious differences in the overall time vigilant (consistent: 7.8 s ± 7.2 s (median ± IQR); inconsistent: 8.1 s ± 6.4 s; full-null model comparison: χ^2^_1_ =0.04, *P*=0.84, table A5). We found no evidence that unit size influenced any of the response variables (Duration: *P*=0.38; Latency: *P*=0.63, Vigilance: *P*=0.15, tables A3, A4, A5). The presence of other parties during the trial did not affect the responses of the males (Duration: *P*=0.94; Latency: *P*=0.73, Vigilance: *P*=0.89, tables A2.7, A2.8, A2.9).

## Discussion

Male Guinea baboons showed no signs of surprise when calls from associated females were played back from an impossible location. Instead, they responded equally strongly to playbacks of calls from an impossible or a possible location. Further, males responded more strongly to the playback of vocalizations from unit-females compared to non-unit-females. While males seemed to be able to recognise their unit’s females by voice, they lacked either the ability or the motivation to track their females’ positions.

These findings were not in line with our initial prediction that primary males monitor the whereabouts of their females. Guinea baboons form one-male units similar to hamadryas baboons or mountain gorillas (*Gorilla b. beringei*). In both of these species, sexual coercion (Smuts & Smuts, 1993) is used by leader males to control female movement and interactions and to prevent transfers to other males (Schreier & Swedell, 2009; Sicotte, 1993). In Guinea baboons, we did not observe such overt aggression towards females, except for some occasional chasing of females. Indeed, female Guinea baboons can roam relatively unimpeded and interact socially with other group members, including other adult males (Goffe et al., 2016).

The lack of differentiated response fits with the relatively laid-back stance of Guinea baboon males. Males form strong bonds with other males (Dal Pesco et al., 2021, 2022; Patzelt et al., 2014). They also show low levels of overt aggression, preventing us from discerning a clear dominance hierarchy (Dal Pesco et al., 2021). At the same time, female Guinea baboons have considerable leverage in mate choice and intersexual bond maintenance (Goffe et al., 2016). Male strategies mainly seem to consist of investing their social time into female grooming and support. Interestingly, males appear to face a trade-off in the allocation of social time, as male investment into socio-positive interactions with other male declines with increasing unit size (Dal Pesco et al., 2022). Social investment into females thus might be important for intersexual bond maintenance and potentially female mate choice in the first place.

Since we tested males when the female whose calls were played was not receptive, we do not know whether males would be more attentive if the female would be able to conceive. We conducted the trials only while females were non-receptive because, during females’ oestrus, primary males and females are less likely to separate (Goffe, 2016), leaving very few opportunities for conducting the experimental trials. Thus, we cannot exclude the possibility that males would respond differentially in conditions where they should be more motivated to track their female’s whereabouts. It is important to note that we do not make any claims about the males’ ability to track female whereabouts – it might well be that they are aware of their females’ locations but simply do not care to attend to apparent violations of their expectations. This inability to distinguish between what the animals ‘can do’ and ‘do do’ is, unfortunately, one of the limitations of such kinds of field experiments (Fischer, 2022).

Our study adds to the accumulating evidence that the need to monitor the social environment varies between species with the degree of competition among individuals. For instance, the highly competitive chacma baboons (*Papio ursinus*), which live in female-philopatric groups, show strong responses to the playback of vocalisation from unfamiliar males (Kitchen et al., 2005, 2013), while Guinea baboons showed greater attention to vocalisations from familiar males compared to neighbours or strangers (Maciej, Patzelt, et al., 2013). In geladas (*Theropithecus gelada*), which live in a multi-level society in aggregations of up to several hundred individuals, vocal recognition seems to be limited to individuals with a high degree of social overlap (Bergman, 2010). Additionally, when presenting individuals with information about changes in association patterns, chacma baboons responded strongly to simulated separations of consortships (Crockford et al., 2007), while Guinea baboons paid more attention to information consistent with current male-female association patterns (Faraut & Fischer, 2019). Similarly, Geladas did not differentiate between consistent or inconsistent information about male-female relationships at all (le Roux & Bergman, 2012).

While the link between group-living and sophisticated social knowledge is well documented (Seyfarth & Cheney, 2015), it is still unclear whether life in a socially complex environment per se (Holekamp, 2007) or rather the degree of competition within and between groups selects for advanced socio-cognitive skills (“Machiavellian intelligence”; Whiten & Byrne, 1988). Deciding on this issue is further complicated by the question of how to operationalize social complexity.

Kappeler (2019) provided a qualitative framework for social complexity, which encompassed Social organization, Social structure, the Mating system, and the Care system. Lukas and Clutton-Brock, 2018 distinguished between organisational complexity in animal societies, referring to the division of labour, and relational complexity, referring to the differentiation of social relationships among group members. They reject the idea of a unitary concept of social complexity (and we agree). Other authors have proposed to hone in on the differentiation of social relationships. Bergman & Beehner (2015) conceived social complexity in terms of the number of differentiated relationships a given individual maintains. Building on this idea, Fischer et al. (2017) proposed a method to quantify social complexity by assessing the diversity of individuals’ relationships. Despite the variety of approaches to operationalising social complexity, the concept, unfortunately, remains elusive. Therefore, the present results cannot more specifically inform the debate between the link between social complexity and social cognition.

With regards to the link between the degree of competition between individuals and the allocation of social attention, more progress has been made. Bergman (2010, p. 2050) argued that “missing social knowledge” might be a consequence of the absence of a competitive environment that offers no benefits for the ability to assess and use of specific social information of conspecifics. Our results, as well as results from previous studies on the same population (Faraut & Fischer, 2019; Maciej, Patzelt, et al., 2013), suggest that a reduced competitive environment affects the value of social information, and as a consequence, the motivation or ability of an individual to attend to them. At the same time, both Guinea baboons and geladas live in highly structured multi-level groups, suggesting that a complex social organisation does not per se select for a high motivation to monitor the social environment. We contend that a skewed distribution of power influences the value of social information and therefore the motivation to attend to events in the social environment.

## Supporting information

Data Analyses

Experiment Data Files

## Acknowledgements

We thank the Diréction des Parcs Nationaux and Ministère de l’Environnement et de la Protection de la Nature du Sénégal for permission to work in the Niokolo-Koba National Park. We are grateful to all research assistants for their help in collecting data.

### Appendix

#### Figures

**Figure A1.**
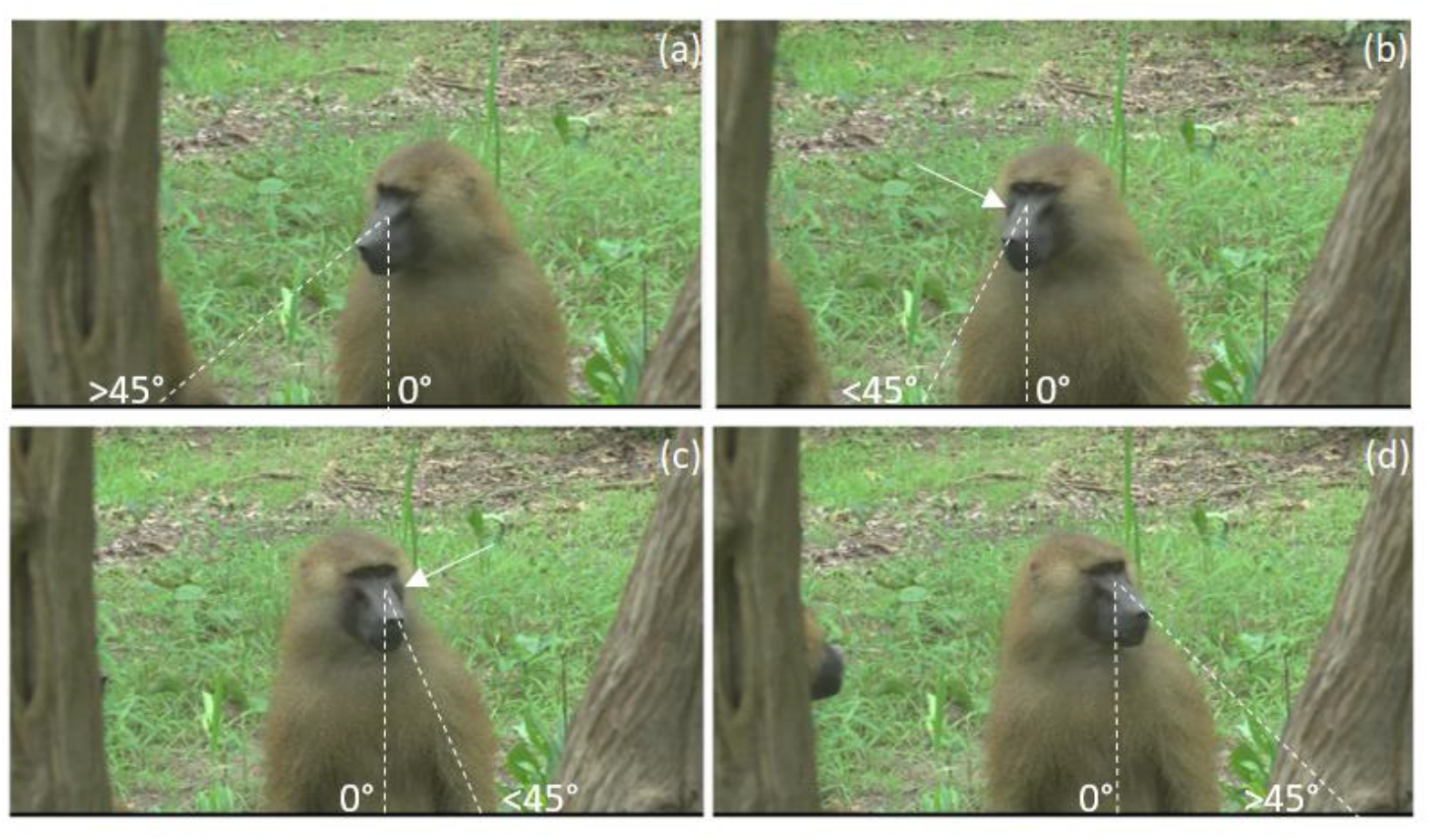
First look duration for males in the a) individual recognition experiment and b) spatial monitoring experiment. Connected points represent data from the same individual (a: *N*=14; b: *N*=22).

#### Tables

**Table A1.** Results of linear mixed model analysing the influence of the main predictor experimental condition (unit-female vs. non-unit-female) and unit size (number of females) on looking duration for experiment 1 (individual recognition).

| Term | Estimate | SE | lower CI | upper CI | t | df | P | min | max |
| --- | --- | --- | --- | --- | --- | --- | --- | --- | --- |
| Intercept | 1.269 | 0.289 | 0.637 | 1.898 | (1) | (1) | (1) | 0.576 | 1.653 |
| Condition | 0.664 | 0.181 | 0.248 | 1.108 | 3.275 | 1 | 0.008 | -0.111<sup>(2)</sup> | 1.198 |
| Nr_Females | -0.075 | 0.081 | -0.258 | 0.109 | -0.669 | 1 | 0.521 | -0.192 | 0.080 |

SE: standard error, CI: confidence interval, min/max: minimum and maximum estimate from model stability analyses. (1) Not presented due to very limited interpretability. (2) Model stability test without dysfunctional model (table A2) leads to different range of estimates (min: 0.478, max: 1.198). Model formula: $\text{lmer}(\log(\text{First\_Look\_Dur}) \sim \text{Condition} + \text{Nr\_Females} + (1|\text{MaleID}) + (1|\text{FemaleIID}) + (1|\text{CallID}))$ .

**Table A2.**
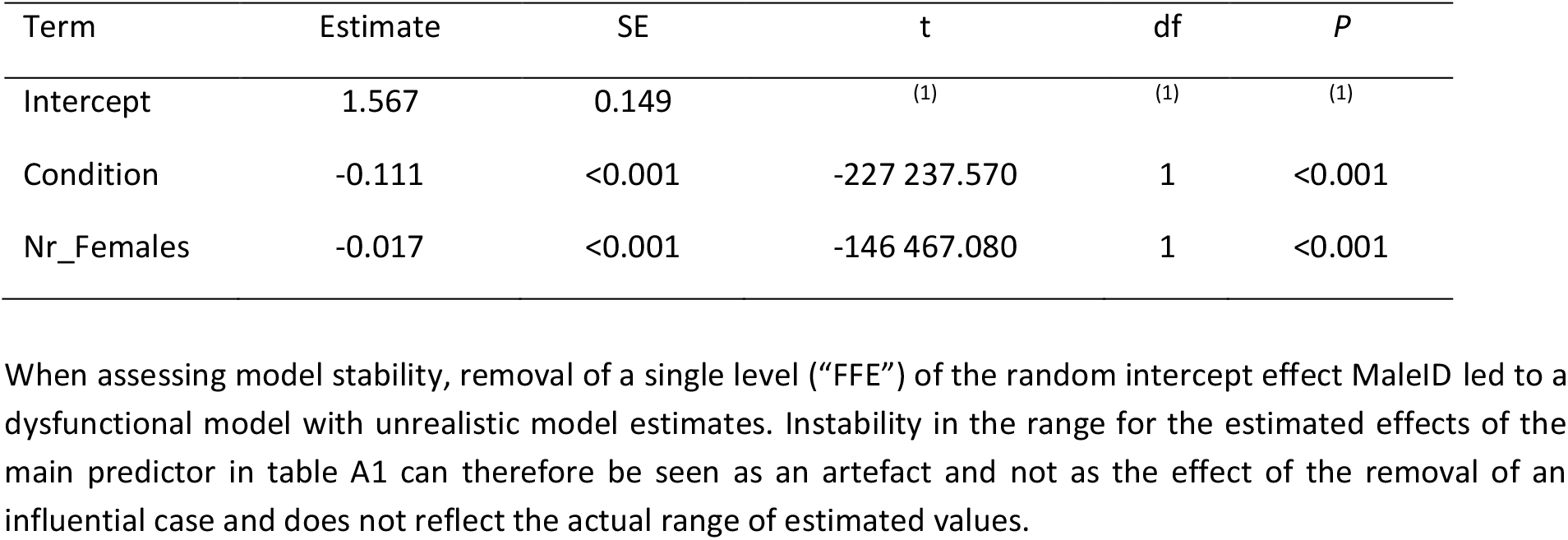
Reduced linear mixed model analysing the influence of the main predictor experimental condition (unit-female vs. non-unit-female) and unit size (number of females) on looking duration for experiment 1 (individual recognition).

| Term | Estimate | SE | t | df | P |
| --- | --- | --- | --- | --- | --- |
| Intercept | 1.567 | 0.149 | (1) | (1) | (1) |
| Condition | -0.111 | <0.001 | -227 237.570 | 1 | <0.001 |
| Nr_Females | -0.017 | <0.001 | -146 467.080 | 1 | <0.001 |

When assessing model stability, removal of a single level ("FFE") of the random intercept effect MaleID led to a dysfunctional model with unrealistic model estimates. Instability in the range for the estimated effects of the main predictor in table A1 can therefore be seen as an artefact and not as the effect of the removal of an influential case and does not reflect the actual range of estimated values.

**Table A3.** Results of Cox proportional hazard model analysing the influence of the main predictor experimental condition (unit-female vs. non-unit-female) and unit size (number of females) on looking latency for experiment 1 (individual recognition).

| Term | Estimate | SE | lower CI | upper CI | Chi<sup>2</sup> | df | P | min | max |
| --- | --- | --- | --- | --- | --- | --- | --- | --- | --- |
| Condition | 0.738 | 2.091 | (1) | (1) | 2.459 | 1 | 0.117 | 0.468 | 0.963 |
| Nr_Females | 0.247 | 1.280 | (1) | (1) | 0.809 | 1 | 0.369 | 0.123 | 0.495 |

SE: standard error, CI: confidence interval, min/max: minimum and maximum estimate from model stability analyses. (1) Confidence interval could not be calculated for Cox proportional hazard model with more than one random effect included. Model formula: `coxme(Surv(First_Look_Lat) ~ Condition + Nr_Females + (1|MaleID) + (1|FemaleID) + (1|CallID))`.

**Table A4.** Results of generalised linear mixed model analysing the influence of the main predictor experimental condition (consistent vs. inconsistent) and unit size (number of females) on looking duration for experiment 2 (spatial monitoring).

| Term | Estimate | SE | lower CI | upper CI | t | df | P | min | max |
| --- | --- | --- | --- | --- | --- | --- | --- | --- | --- |
| Intercept | 1.529 | 0.469 | 0.739 | 2.068 | (1) | (1) | (1) | 1.212 | 1.661 |
| Condition | -0.005 | 0.341 | -0.538 | 0.517 | <0.001 | 1 | 0.989 | -0.079 | 0.142 |
| Nr_Females | -0.145 | 0.165 | -0.348 | 0.086 | -0.669 | 1 | 0.381 | -0.259 | 0.018 |

SE: standard error, CI: confidence interval, min/max: minimum and maximum estimate from model stability analyses. (1) Not presented due to very limited interpretability. Model formula: `glmer(tr.First_Look_Dur ~ Condition + Nr_Females + (1|MaleID), family = Gamma(link = "log"))`.

**Table A5.** Results of Cox proportional hazard model analysing the influence of the main predictor experimental condition (consistent vs. inconsistent) and unit size (number of females) on looking latency for experiment 2 (spatial monitoring).

| Term | Estimate | SE | lower CI | upper CI | Chi<sup>2</sup> | df | P | min | max |
| --- | --- | --- | --- | --- | --- | --- | --- | --- | --- |
| Condition | 0.318 | 1.374 | -0.246 | 1.039 | 1.104 | 1 | 0.293 | 0.150 | 1.466 |
| Nr_Females | -0.097 | 0.908 | -0.556 | 0.563 | 0.232 | 1 | 0.630 | -0.178 | 0.109 |

SE: standard error, CI: confidence interval, min/max: minimum and maximum estimate from model stability analyses. Model formula: `coxme(Surv(First_Look_Lat) ~ Condition + Nr_Females + (1|MaleID))`.

**Table A6:** Results of linear mixed model analysing the influence of the main predictor experimental condition (consistent vs. inconsistent) and unit size (number of females) on vigilance time for experiment 2 (spatial monitoring).

| Term | Estimate | SE | lower CI | upper CI | t | df | P | min | max |
| --- | --- | --- | --- | --- | --- | --- | --- | --- | --- |
| Intercept | 3.335 | 0.380 | 2.600 | 4.120 | (1) | (1) | (1) | 2.986 | 3.634 |
| Condition | 0.061 | 0.299 | -0.530 | 0.660 | 0.202 | 1 | 0.841 | -0.123 | 0.167 |
| Nr_Females | -0.207 | 0.125 | -0.460 | 0.050 | -1.509 | 1 | 0.148 | -0.284 | -0.047 |

SE: standard error, CI: confidence interval, min/max: minimum and maximum estimate from model stability analyses. (1) Not presented due to very limited interpretability. Model formula: `lmer(Vigilance2 ~ Condition + Nr_Females + (1|MaleID))`.

**Table A7.** Model estimates for control predictor Mingled of alternative generalized linear mixed model (compare table A3; First-Look Duration) for experiment 2 (spatial monitoring) including the predictor Mingled (Y/N) to indicate the presence of another baboon party during the trial.

| Term | Estimate | SE | lower CI | upper CI | t | df | P | min | max |
| --- | --- | --- | --- | --- | --- | --- | --- | --- | --- |
| Intercept | 1.505 | 0.569 | 0.540 | 2.251 | (1) | (1) | (1) | 1.220 | 1.621 |
| Condition | 0.002 | 0.354 | -0.545 | 0.551 | 0.007 | 1 | 0.994 | -0.079 | 0.224 |
| Nr_Females | -0.143 | 0.167 | -0.362 | 0.072 | -0.852 | 1 | 0.394 | -0.256 | 0.017 |
| Mingled | 0.028 | 0.361 | -0.555 | 0.641 | 0.077 | 1 | 0.939 | -0.065 | 0.216 |

SE: standard error, CI: confidence interval, min/max: minimum and maximum estimate from model stability analyses. (1) Not presented due to very limited interpretability. Model formula: `glmer(tr.First_Look_Dur~Condition + Nr_Females + Mingled + (1|MaleID), family = Gamma(link = "log"))`.

**Table A8.**
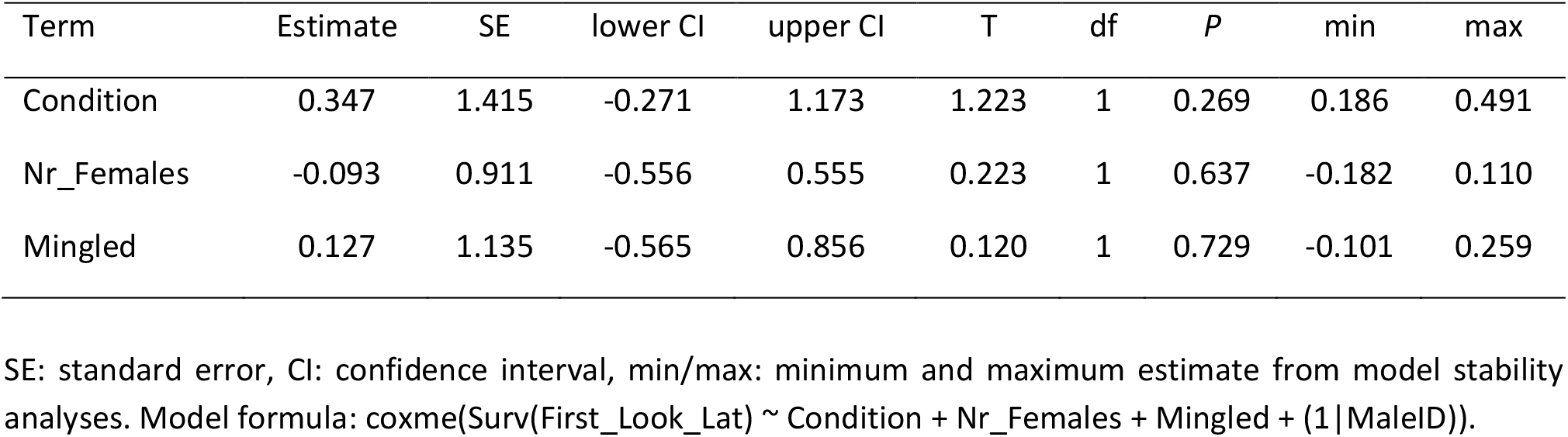
Model estimates for control predictor *Mingled* of alternative Cox proportional hazard model (compare table A4, First-Look Latency) for experiment 2 (spatial monitoring) including the predictor *Mingled* (Y/N) to indicate the presence of another baboon party during the trial.

**Table A9:** Model estimates for control predictor *Mingled* of alternative linear mixed model (compare table A5, Vigilance Time) for experiment 2 (spatial monitoring) including the predictor *Mingled* (Y/N) to indicate the presence of another baboon party during the trial.

| Term | Estimate | SE | lower CI | upper CI | t | df | P | min | max |
| --- | --- | --- | --- | --- | --- | --- | --- | --- | --- |
| Intercept | 3.372 | 0.470 | 2.394 | 4.331 | (1) | (1) | (1) | 2.991 | 3.914 |
| Condition | 0.047 | 0.318 | -0.532 | 0.663 | 0.142 | 1 | 0.888 | -0.139 | 0.195 |
| Nr_Females | -0.209 | 0.127 | -0.469 | 0.039 | -1.498 | 1 | 0.150 | -0.308 | -0.047 |
| Mingled | -0.045 | 0.329 | -0.671 | 0.605 | -0.138 | 1 | 0.891 | -0.301 | 0.124 |

SE: standard error, CI: confidence interval, min/max: minimum and maximum estimate from model stability analyses. (1) Not presented due to very limited interpretability. Model formula: `lmer(Vigilance2 ~ Condition + Nr_Females + Mingled + (1|MaleID))`.

## References

Baayen, R. H., Davidson, D. J., & Bates, D. M. (2008). Mixed-effects modeling with crossed random effects for subjects and items. Journal of Memory and Language, 59(4), 390–412. https://doi.org/10.1016/j.jml.2007.12.005

Balsby, T. J., & Dabelsteen, T. (2005). Simulated courtship interactions elicit neighbour intrusions in the whitethroat, Sylvia communis. Animal Behaviour, 69(1), 161–168. https://doi.org/10.1016/j.anbehav.2004.01.021

Bergman, T. J. (2010). Experimental evidence for limited vocal recognition in a wild primate: Implications for the social complexity hypothesis. Proceedings of the Royal Society B: Biological Sciences, 277(1696), 3045–3053. https://doi.org/10.1098/rspb.2010.0580

Bergman, T. J., & Beehner, J. C. (2015). Measuring social complexity. Animal Behaviour, 103, 203–209. https://doi.org/10.1016/j.anbehav.2015.02.018

Birkhead, T. R., & Møller, A. P. (1995). Extra-pair copulation and extra-pair paternity in birds. Animal Behaviour, 49(3), 843–848.

Clutton-Brock, T. H., & Parker, G. A. (1992). Potential Reproductive Rates and the Operation of Sexual Selection. The Quarterly Review of Biology, 67(4), 437–456. https://doi.org/10.1086/417793

Clutton-Brock, T. H., & Vincent, A. C. J. (1991). Sexual selection and the potential reproductive rates of males and females. Nature, 351(6321), 58–60. https://doi.org/10.1038/351058a0

Crockford, C., Wittig, R. M., Seyfarth, R. M., & Cheney, D. L. (2007). Baboons eavesdrop to deduce mating opportunities. Animal Behaviour, 73(5), 885–890. https://doi.org/10.1016/j.anbehav.2006.10.016

Dal Pesco, F., Trede, F., Zinner, D., & Fischer, J. (2021). Kin bias and male pair-bond status shape male-male relationships in a multilevel primate society. Behavioral Ecology and Sociobiology, 75(1), 1–14. https://doi.org/10.1007/s00265-020-02960-8

Dal Pesco, F., Trede, F., Zinner, D., & Fischer, J. (2022). Male–male social bonding, coalitionary support and reproductive success in wild Guinea baboons. Proceedings of the Royal Society B, 289(1975), 20220347. https://doi.org/10.1098/rspb.2022.0347

Davies, A. D., Lewis, Z., & Dougherty, L. R. (2020). A meta-analysis of factors influencing the strength of mate-choice copying in animals. Behavioral Ecology, 31(6), 1279–1290. https://doi.org/10.1093/beheco/araa064

Dobson, A. J., & Barnett, A. G. (2018). An introduction to generalized linear models. Chapman and Hall/CRC.

Engh, A. L., Siebert, E. R., Greenberg, D. A., & Holekamp, K. E. (2005). Patterns of alliance formation and postconflict aggression indicate spotted hyaenas recognize third-party relationships. Animal Behaviour, 69(1), 209–217. https://doi.org/10.1016/j.anbehav.2004.04.013

Faraut, L., & Fischer, J. (2019). How life in a tolerant society affects the attention to social information in baboons. Animal Behaviour, 152, 11–17. https://doi.org/10.1016/j.anbehav.2019.04.004

Fischer, J. (2022). Studying primate cognition: From the wild to captivity and back. Primate Cognitive Studies; Schwartz, B., Beran, M., Eds, 609–631.

Fischer, J., Farnworth, M. S., Sennhenn-Reulen, H., & Hammerschmidt, K. (2017). Quantifying social complexity. Animal Behaviour, 130, 57–66.

Fischer, J., Kopp, G. H., Dal Pesco, F., Goffe, A., Hammerschmidt, K., Kalbitzer, U., Klapproth, M., Maciej, P., Ndao, I., Patzelt, A., & Zinner, D. (2017). Charting the neglected West: The social system of Guinea baboons. American Journal of Physical Anthropology, 162(S63), 15–31. https://doi.org/10.1002/ajpa.23144

Fischer, J., Noser, R., & Hammerschmidt, K. (2013). Bioacoustic Field Research: A Primer to Acoustic Analyses and Playback Experiments With Primates: Bioacoustic Field Methods. American Journal of Primatology, 75(7), 643–663. https://doi.org/10.1002/ajp.22153

Goffe, A. S. (2016). Social relationships of female Guinea baboons (Papio papio) in Senegal. Dissertation, Göttingen, Georg-August Universität, 2016.

Goffe, A. S., Zinner, D., & Fischer, J. (2016). Sex and friendship in a multilevel society: Behavioural patterns and associations between female and male Guinea baboons. Behavioral Ecology and Sociobiology, 70(3), 323–336. https://doi.org/10.1007/s00265-015-2050-6

Guidelines for the treatment of animals in behavioural research and teaching. (2020). Animal Behaviour, 159, I–XI. https://doi.org/10.1016/j.anbehav.2019.11.002

Holekamp, K. E. (2007). Questioning the social intelligence hypothesis. Trends in Cognitive Sciences, 11(2), 65–69. https://doi.org/10.1016/j.tics.2006.11.003

Jahn-Eimermacher, A., Lasarzik, I., & Raber, J. (2011). Statistical analysis of latency outcomes in behavioral experiments. Behavioural Brain Research, 221(1), 271–275. https://doi.org/10.1016/j.bbr.2011.03.007

Kappeler, P. M., Clutton-Brock, T., Shultz, S., & Lukas, D. (2019). Social complexity: Patterns, processes, and evolution. Behavioral Ecology and Sociobiology, 73(1), 5, s00265-018-2613– 2614. https://doi.org/10.1007/s00265-018-2613-4

Kitchen, D. M., Cheney, D. L., Engh, A. L., Fischer, J., Moscovice, L. R., & Seyfarth, R. M. (2013). Male baboon responses to experimental manipulations of loud “wahoo calls”: Testing an honest signal of fighting ability. Behavioral Ecology and Sociobiology, 67(11), 1825–1835. https://doi.org/10.1007/s00265-013-1592-8

Kitchen, D. M., Cheney, D. L., & Seyfarth, R. M. (2005). Male chacma baboons (Papio hamadryas ursinus) discriminate loud call contests between rivals of different relative ranks. Animal Cognition, 8(1), 1–6. https://doi.org/10.1007/s10071-004-0222-2

le Roux, A., & Bergman, T. J. (2012). Indirect rival assessment in a social primate, Theropithecus gelada. Animal Behaviour, 83(1), 249–255. https://doi.org/10.1016/j.anbehav.2011.10.034

Lukas, D., & Clutton-Brock, T. (2018). Social complexity and kinship in animal societies. Ecology Letters, 21(8), 1129–1134. https://doi.org/10.1111/ele.13079

Maciej, P., Fischer, J., & Hammerschmidt, K. (2011). Transmission characteristics of primate vocalizations: Implications for acoustic analyses. PloS One, 6(8), e23015. https://doi.org/10.1371/journal.pone.0023015

Maciej, P., Ndao, I., Hammerschmidt, K., & Fischer, J. (2013). Vocal communication in a complex multi-level society: Constrained acoustic structure and flexible call usage in Guinea baboons. Frontiers in Zoology, 10(1), 58. https://doi.org/10.1186/1742-9994-10-58

Maciej, P., Patzelt, A., Ndao, I., Hammerschmidt, K., & Fischer, J. (2013). Social monitoring in a multilevel society: A playback study with male Guinea baboons. Behavioral Ecology and Sociobiology, 67(1), 61–68. https://doi.org/10.1007/s00265-012-1425-1

Manser, M. B., Allen, C., & Townsend, S. W. (2011). A simple test of vocal individual recognition in wild meerkats. https://doi.org/10.1098/rsbl.2011.0844

Patzelt, A., Kopp, G. H., Ndao, I., Kalbitzer, U., Zinner, D., & Fischer, J. (2014). Male tolerance and male–male bonds in a multilevel primate society. Proceedings of the National Academy of Sciences, 111(41), 14740–14745. https://doi.org/10.1073/pnas.1405811111

Paz-y-MiñoC, G., Bond, A. B., Kamil, A. C., & Balda, R. P. (2004). Pinyon jays use transitive inference to predict social dominance. Nature, 430(7001), Article 7001. https://doi.org/10.1038/nature02723

R Core Team. (2021). R: A Language and Environment for Statistical Computing. R Foundation for Statistical Computing. URL https://www.R-project.org/

Rubenstein, D. I., & Hack, M. (2004). Natural and sexual selection and the evolution of multi-level societies: Insights from zebras with comparisons to primates. In P. M. Kappeler & C. P. van Schaik (Eds.), Sexual Selection in Primates (1st ed., pp. 266–279). Cambridge University Press. https://doi.org/10.1017/CBO9780511542459.017

Schreier, A. L., & Swedell, L. (2009). The fourth level of social structure in a multi-level society: Ecological and social functions of clans in hamadryas baboons. American Journal of Primatology, 71(11), 948–955. https://doi.org/10.1002/ajp.20736

Seyfarth, R. M., & Cheney, D. L. (2015). Social cognition. Animal Behaviour, 103, 191–202. https://doi.org/10.1016/j.anbehav.2015.01.030

Sicotte, P. (1993). Inter-group encounters and female transfer in mountain gorillas: Influence of group composition on male behavior. American Journal of Primatology, 30(1), 21–36. https://doi.org/10.1002/ajp.1350300103

Smuts, B. B., & Smuts, R. W. (1993). Male aggression and sexual coercion of females in nonhuman primates and other mammals: Evidence and theoretical implications. In Advances in the Study of Behavior (Vol. 22, pp. 1–63).

Swedell, L., & Plummer, T. (2012). A Papionin Multilevel Society as a Model for Hominin Social Evolution. International Journal of Primatology, 33(5), 1165–1193. https://doi.org/10.1007/s10764-012-9600-9

Whitehouse, J., & Meunier, H. (2020). An understanding of third-party friendships in a tolerant macaque. Scientific Reports, 10(1), 9777. https://doi.org/10.1038/s41598-020-66407-w

Whiten, A., & Byrne, R. W. (1988). The Machiavellian intelligence hypotheses. Clarendon Press/Oxford University Press.

Wiley, R. H. (2013). Specificity and multiplicity in the recognition of individuals: Implications for the evolution of social behaviour. Biological Reviews, 88(1), 179–195. https://doi.org/10.1111/j.1469-185X.2012.00246.x

