## Supplementary material for "Male Guinea baboons are oblivious to their females’ whereabouts": Data Analyses: boot_glmm.r

```
boot.glmm.pred<-function(model.res, excl.warnings=F, nboots=1000, para=F, resol=1000, level=0.95, use=NULL, circ.var.name=NULL, circ.var=NULL, use.u=F, 
	n.cores=c("all-1", "all"), save.path=NULL, load.lib=T, lib.loc=.libPaths(), set.all.effects.2.zero=F){
	if(load.lib){library(lme4, lib.loc=lib.loc)}
	n.cores=n.cores[1]
	keepWarnings<-function(expr){
		localWarnings <- list()
		value <- withCallingHandlers(expr,
			warning = function(w) {
				localWarnings[[length(localWarnings)+1]] <<- w
				invokeRestart("muffleWarning")
			}
		)
		list(value=value, warnings=localWarnings)
	}
	##define function extracting all estimated coefficients (fixed and random effects) and also the model summary wrt random effects:
	extract.all<-function(mres){
		##extract random effects model summary:
		vc.mat=as.data.frame(summary(mres)$varcor)##... and prepare/extract variance covariance matrix from the model handed over
		xx=lapply(summary(mres)$varcor, function(x){attr(x, "stddev")})##append residual variance
		##create vector with names of the terms in the model
		xnames=c(names(fixef(mres)), paste(rep(names(xx), unlist(lapply(xx, length))), unlist(lapply(xx, names)), sep="@"))
		if(class(mres)[1]=="lmerMod"){xnames=c(xnames, "Residual")}##append "Residual" to xnames in case of Gaussian model
		if(class(mres)[1]=="lmerMod"){res.sd=vc.mat[vc.mat$grp=="Residual", "sdcor"]}##extract residual sd i case of Gaussian model
		#if(vc.mat$grp[nrow(vc.mat)]=="Residual"){vc.mat=vc.mat[-nrow(vc.mat), ]}##and drop residuals-row from vc.mat in case of Gaussisn model
		##deal with names in vc.mat which aren't exactly the name of the resp. random effect
		r.icpt.names=names(ranef(mres))##extract names of random intercepts...
		not.r.icpt.names=setdiff(vc.mat$grp, r.icpt.names)##... and names of the random effects having random slopes
		not.r.icpt.names=unlist(lapply(strsplit(not.r.icpt.names, split=".", fixed=T), function(x){paste(x[1:(length(x)-1)], collapse=".")}))
		vc.mat$grp[!vc.mat$grp%in%r.icpt.names]=not.r.icpt.names
		if(vc.mat$grp[nrow(vc.mat)]=="Residual"){vc.mat$var1[nrow(vc.mat)]=""}
		xnames=paste(vc.mat$grp, vc.mat$var1, sep="@")
		re.summary=unlist(vc.mat$sdcor)
		names(re.summary)=xnames
		ranef(mres)
		##extract random effects model summary: done
		##extract random effects details:
		re.detail=ranef(mres)
		xnames=paste(
			rep(x=names(re.detail), times=unlist(lapply(re.detail, function(x){nrow(x)*ncol(x)}))),
				unlist(lapply(re.detail, function(x){rep(colnames(x), each=nrow(x))})), 
				unlist(lapply(re.detail, function(x){rep(rownames(x), times=ncol(x))})),
				sep="@")
		re.detail=unlist(lapply(re.detail, function(x){
			return(unlist(c(x)))
		}))
		#browser()
		#deal with negative binomial model to extract theta:
		xx=as.character(summary(mres)$call)
		if(any(grepl(x=xx, pattern="negative.binomial"))){
			xx=xx[grepl(x=xx, pattern="negative.binomial")]
			xx=gsub(x=xx, pattern="negative.binomial(theta = ", replacement="", fixed=T)
			xx=gsub(x=xx, pattern=")", replacement="", fixed=T)
			re.detail=c(re.detail, as.numeric(xx))
			xnames=c(xnames, "theta")
		}
		names(re.detail)=xnames
		ns=c(length(fixef(mres)), length(re.summary), length(re.detail))
		names(ns)=c("n.fixef", "n.re.summary", "n.re.detail")
		return(c(fixef(mres), re.summary, re.detail, ns))
	}	
	if(excl.warnings){
		boot.fun<-function(x, model.res., keepWarnings., use.u., save.path.){
			xdone=F
			while(!xdone){
				i.res=keepWarnings(bootMer(x=model.res., FUN=extract.all, nsim=1, use.u=use.u.)$t)
				if(length(unlist(i.res$warnings)$message)==0){
					xdone=T
				}
			}
			est.effects=i.res$value
			i.warnings=NULL
			if(length(save.path.)>0){save(file=paste(c(save.path., "/b_", x, ".RData"), collapse=""), list=c("est.effects", "i.warnings"))}
			return(i.res$value)
		}
	}else{
		boot.fun<-function(y, model.res., keepWarnings., use.u., save.path.){
			#keepWarnings.(bootMer(x=model.res., FUN=fixef, nsim=1)$t)
			i.res=keepWarnings(bootMer(x=model.res., FUN=extract.all, nsim=1, use.u=use.u.))
			if(length(save.path.)>0){
				est.effects=i.res$value$t
				i.warnings=i.res$warnings
				save(file=paste(c(save.path., "/b_", y, ".RData"), collapse=""), list=c("est.effects", "i.warnings"))
			}
			return(list(ests=i.res$value$t, warns=unlist(i.res$warnings)))
		}
	}
	if(para){
		on.exit(expr = parLapply(cl=cl, X=1:length(cl), fun=function(x){rm(list=ls())}), add = FALSE)
		on.exit(expr = stopCluster(cl), add = T)
		library(parallel)
		cl <- makeCluster(getOption("cl.cores", detectCores()))
		if(n.cores!="all"){
			if(n.cores=="all-1"){n.cores=length(cl)-1}
			if(n.cores<length(cl)){
				cl=cl[1:n.cores]
			}
		}
		parLapply(cl=cl, 1:length(cl), fun=function(x, .lib.loc=lib.loc){
		  library(lme4, lib.loc=.lib.loc)
		  return(invisible(""))
		})
		all.res=parLapply(cl=cl, X=1:nboots, fun=boot.fun, model.res.=model.res, keepWarnings.=keepWarnings, use.u.=use.u, save.path.=save.path)
	}else{
    all.res=lapply(X=1:nboots, FUN=boot.fun, model.res.=model.res, use.u.=use.u, save.path.=save.path)#, keepWarnings=excl.warnings)
	}
	if(!excl.warnings){
		all.warns=lapply(all.res, function(x){
			xxx=unlist(x$warns)
			if(length(xxx)==0){xxx=""}
			return(xxx)
		})
		all.res=lapply(all.res, function(x){x$ests})
	}else{
		all.warns=NULL
	}
	if(length(use)>0){
		#extract fixed effects terms from the model:
		xcall=as.character(model.res@call)[2]
		model.terms=attr(terms(as.formula(xcall)), "term.labels")
		REs=names(ranef(model.res))
		#for(i in 1:length(REs)){
			model.terms=model.terms[!grepl(x=model.terms, pattern="|", fixed=T)]
		#}
		model=paste(model.terms, collapse="+")
		#build model wrt to the fixed effects:
		model.terms=unique(unlist(strsplit(x=model.terms, split=":", fixed=T)))
		model.terms=model.terms[!grepl(x=model.terms, pattern="I(", fixed=T)]
		model.terms=model.terms[!grepl(x=model.terms, pattern="^2)", fixed=T)]
		#exclude interactions and squared terms from model.terms:
		#create new data to be used to determine fitted values:
		ii.data=model.res@frame
		
		if(length(circ.var.name)==1){
			set.circ.var.to.zero=sum(circ.var.name%in%use)==0
		}else{
			set.circ.var.to.zero=F
		}
		
		if(length(use)==0){use=model.terms}
		new.data=vector("list", length(model.terms))
		usel=model.terms%in%use
		#if(length(use)>0)
		for(i in 1:length(model.terms)){
			if(is.factor(ii.data[, model.terms[i]])){
				new.data[[i]]=levels(ii.data[, model.terms[i]])
			}else if(!is.factor(ii.data[, model.terms[i]]) & usel[i] & ifelse(length(circ.var.name)==0, T, !grepl(x=model.terms[i], pattern=circ.var.name))){
				new.data[[i]]=seq(from=min(ii.data[, model.terms[i]]), to=max(ii.data[, model.terms[i]]), length.out=resol)
			}else  if(!is.factor(ii.data[, model.terms[i]]) & ifelse(length(circ.var.name)==0, T, !grepl(x=model.terms[i], pattern=circ.var.name))){
				new.data[[i]]=mean(ii.data[, model.terms[i]])
			}
		}
		names(new.data)=model.terms
		if(length(circ.var.name)==1){
			new.data=new.data[!(model.terms%in%paste(c("sin(", "cos("), circ.var.name, ")", sep=""))]
			if(sum(grepl(pattern=circ.var.name, x=use))>0){
				new.data=c(new.data, list(seq(min(circ.var, na.rm=T), max(circ.var, na.rm=T), length.out=resol)))
				names(new.data)[length(new.data)]=circ.var.name
			}else{
				new.data=c(new.data, list(0))
			}
			model.terms=model.terms[!(model.terms%in%paste(c("sin(", "cos("), circ.var.name, ")", sep=""))]
		}
		xnames=names(new.data)
		#browser()
		new.data=data.frame(expand.grid(new.data))
		names(new.data)=xnames
		#names(new.data)[1:length(model.terms)]=model.terms
		#browser()
		if(length(circ.var.name)==1){
			names(new.data)[ncol(new.data)]=circ.var.name
		}
		#create predictors matrix:
# 		if(length(circ.var.name)>0 & length(intersect(circ.var.name, names(new.data)))>0){
# 			new.data=cbind(new.data, sin(new.data[, circ.var.name]), cos(new.data[, circ.var.name]))
# 			names(new.data)[(ncol(new.data)-1):ncol(new.data)]=paste(c("sin(", "cos("), circ.var.name, ")", sep="")
# 			new.data=new.data[, !names(new.data)==circ.var.name]
# 		}
# 		browser()
		r=runif(nrow(new.data))
		if(set.all.effects.2.zero){
			for(iterm in setdiff(colnames(new.data), c("(Intercept)", use))){
				new.data[, iterm]=0
			}
		}
		m.mat=model.matrix(object=as.formula(paste(c("r", model), collapse="~")), data=new.data)
		if(set.circ.var.to.zero){
			m.mat[,paste(c("sin(", circ.var.name, ")"), collapse="")]=0
			m.mat[,paste(c("cos(", circ.var.name, ")"), collapse="")]=0
		}
		#m.mat=t(m.mat)
		#get the CIs for the fitted values:
		ci=lapply(all.res, function(x){
			#return(apply(m.mat[names(fixef(model.res)), , drop=F]*as.vector(x[, names(fixef(model.res))]), 2, sum))
			return(m.mat[, names(fixef(model.res)), drop=F]%*%as.vector(x[, names(fixef(model.res))]))
		})
		ci=matrix(unlist(ci), ncol=nboots, byrow=F)
		ci=t(apply(ci, 1, quantile, prob=c((1-level)/2, 1-(1-level)/2), na.rm=T))
		colnames(ci)=c("lower.cl", "upper.cl")
		fv=m.mat[, names(fixef(model.res))]%*%fixef(model.res)
		if(class(model.res)[[1]]!="lmerMod"){
			if(model.res@resp$family$family=="binomial"){
				ci=exp(ci)/(1+exp(ci))
				fv=exp(fv)/(1+exp(fv))
			}else if(model.res@resp$family$family=="poisson" | (model.res@resp$family$family=="Gamma" & model.res@resp$family$link=="log") | substr(x=model.res@resp$family$family, start=1, stop=17)=="Negative Binomial"){
				ci=exp(ci)
				fv=exp(fv)
			}
		}
		result=data.frame(new.data, fitted=fv, ci)
	}else{
		result=NULL
	}
	all.boots=matrix(unlist(all.res), nrow=nboots, byrow=T)
	colnames(all.boots)=colnames(all.res[[1]])
	ci.est=apply(all.boots, 2, quantile, prob=c((1-level)/2, 1-(1-level)/2), na.rm=T)
	if(length(fixef(model.res))>1){
		ci.est=data.frame(orig=fixef(model.res), t(ci.est)[1:length(fixef(model.res)), ])
	}else{
		ci.est=data.frame(orig=fixef(model.res), t(t(ci.est)[1:length(fixef(model.res)), ]))
	}
	return(list(ci.predicted=result, ci.estimates=ci.est, all.warns=all.warns, all.boots=all.boots))
}
###########################################################################################################
###########################################################################################################
boot.lmer<-function(model.res, excl.warnings=F, nboots=1000, para=F, resol=1000, level=0.95, 
	use=NULL, circ.var.name=NULL, circ.var=NULL, use.u=F, 
	n.cores=c("all-1", "all"), save.path=NULL, load.lib=T, lib.loc=.libPaths(), set.all.effects.2.zero=F){
	if(load.lib){library(lme4, lib.loc=lib.loc)}
	n.cores=n.cores[1]
	if(is.list(use)){use.list=use}else{use.list=list(use)}
	keepWarnings<-function(expr){
		localWarnings <- list()
		value <- withCallingHandlers(expr,
			warning = function(w) {
				localWarnings[[length(localWarnings)+1]] <<- w
				invokeRestart("muffleWarning")
			}
		)
		list(value=value, warnings=localWarnings)
	}
	##define function extracting all estimated coefficients (fixed and random effects) and also the model summary wrt random effects:
	extract.all<-function(mres){
		##extract random effects model summary:
		vc.mat=as.data.frame(summary(mres)$varcor)##... and prepare/extract variance covariance matrix from the model handed over
		xx=lapply(summary(mres)$varcor, function(x){attr(x, "stddev")})##append residual variance
		##create vector with names of the terms in the model
		xnames=c(names(fixef(mres)), paste(rep(names(xx), unlist(lapply(xx, length))), unlist(lapply(xx, names)), sep="@"))
		if(class(mres)[1]=="lmerMod"){xnames=c(xnames, "Residual")}##append "Residual" to xnames in case of Gaussian model
		if(class(mres)[1]=="lmerMod"){res.sd=vc.mat[vc.mat$grp=="Residual", "sdcor"]}##extract residual sd i case of Gaussian model
		#if(vc.mat$grp[nrow(vc.mat)]=="Residual"){vc.mat=vc.mat[-nrow(vc.mat), ]}##and drop residuals-row from vc.mat in case of Gaussisn model
		##deal with names in vc.mat which aren't exactly the name of the resp. random effect
		r.icpt.names=names(ranef(mres))##extract names of random intercepts...
		not.r.icpt.names=setdiff(vc.mat$grp, r.icpt.names)##... and names of the random effects having random slopes
		not.r.icpt.names=unlist(lapply(strsplit(not.r.icpt.names, split=".", fixed=T), function(x){paste(x[1:(length(x)-1)], collapse=".")}))
		vc.mat$grp[!vc.mat$grp%in%r.icpt.names]=not.r.icpt.names
		if(vc.mat$grp[nrow(vc.mat)]=="Residual"){vc.mat$var1[nrow(vc.mat)]=""}
		xnames=paste(vc.mat$grp, vc.mat$var1, sep="@")
		re.summary=unlist(vc.mat$sdcor)
		names(re.summary)=xnames
		ranef(mres)
		##extract random effects model summary: done
		##extract random effects details:
		re.detail=ranef(mres)
		xnames=paste(
			rep(x=names(re.detail), times=unlist(lapply(re.detail, function(x){nrow(x)*ncol(x)}))),
				unlist(lapply(re.detail, function(x){rep(colnames(x), each=nrow(x))})), 
				unlist(lapply(re.detail, function(x){rep(rownames(x), times=ncol(x))})),
				sep="@")
		re.detail=unlist(lapply(re.detail, function(x){
			return(unlist(c(x)))
		}))
		#deal with negative binomial model to extract theta:
		xx=as.character(summary(mres)$call)
		if(any(grepl(x=xx, pattern="negative.binomial"))){
			xx=xx[grepl(x=xx, pattern="negative.binomial")]
			xx=gsub(x=xx, pattern="negative.binomial(theta = ", replacement="", fixed=T)
			xx=gsub(x=xx, pattern=")", replacement="", fixed=T)
			re.detail=c(re.detail, as.numeric(xx))
			xnames=c(xnames, "theta")
		}
		names(re.detail)=xnames
		ns=c(length(fixef(mres)), length(re.summary), length(re.detail))
		names(ns)=c("n.fixef", "n.re.summary", "n.re.detail")
		return(c(fixef(mres), re.summary, re.detail, ns))
	}	
	if(excl.warnings){
		boot.fun<-function(x, model.res., keepWarnings., use.u., save.path.){
			xdone=F
			while(!xdone){
				i.res=keepWarnings(bootMer(x=model.res., FUN=extract.all, nsim=1, use.u=use.u.)$t)
				if(length(unlist(i.res$warnings)$message)==0){
					xdone=T
				}
			}
			est.effects=i.res$value
			i.warnings=NULL
			if(length(save.path.)>0){save(file=paste(c(save.path., "/b_", x, ".RData"), collapse=""), list=c("est.effects", "i.warnings"))}
			return(i.res$value)
		}
	}else{
		boot.fun<-function(y, model.res., keepWarnings., use.u., save.path.){
			#keepWarnings.(bootMer(x=model.res., FUN=fixef, nsim=1)$t)
			i.res=keepWarnings(bootMer(x=model.res., FUN=extract.all, nsim=1, use.u=use.u.))
			if(length(save.path.)>0){
				est.effects=i.res$value$t
				i.warnings=i.res$warnings
				save(file=paste(c(save.path., "/b_", y, ".RData"), collapse=""), list=c("est.effects", "i.warnings"))
			}
			return(list(ests=i.res$value$t, warns=unlist(i.res$warnings)))
		}
	}
	if(para){
		on.exit(expr = parLapply(cl=cl, X=1:length(cl), fun=function(x){rm(list=ls())}), add = FALSE)
		on.exit(expr = stopCluster(cl), add = T)
		library(parallel)
		cl <- makeCluster(getOption("cl.cores", detectCores()))
		if(n.cores!="all"){
			if(n.cores!="all-1"){n.cores=length(cl)-1}
			if(n.cores<length(cl)){
				cl=cl[1:n.cores]
			}
		}
		parLapply(cl=cl, 1:length(cl), fun=function(x, .lib.loc=lib.loc){
		  library(lme4, lib.loc=.lib.loc)
		  return(invisible(""))
		})
		all.res=parLapply(cl=cl, X=1:nboots, fun=boot.fun, model.res.=model.res, keepWarnings.=keepWarnings, use.u.=use.u, save.path.=save.path)
	}else{
    all.res=lapply(X=1:nboots, FUN=boot.fun, model.res.=model.res, use.u.=use.u, save.path.=save.path)#, keepWarnings=excl.warnings)
	}
	if(!excl.warnings){
		all.warns=lapply(all.res, function(x){
			xxx=unlist(x$warns)
			if(length(xxx)==0){xxx=""}
			return(xxx)
		})
		all.res=lapply(all.res, function(x){x$ests})
	}else{
		all.warns=NULL
	}
	if(length(use.list)>0){
		#extract fixed effects terms from the model:
		xcall=as.character(model.res@call)[2]
		model.terms=attr(terms(as.formula(xcall)), "term.labels")
		REs=names(ranef(model.res))
		model.terms=model.terms[!grepl(x=model.terms, pattern="|", fixed=T)]
		model=paste(model.terms, collapse="+")
		#build model wrt to the fixed effects:
		model.terms=unique(unlist(strsplit(x=model.terms, split=":", fixed=T)))
		model.terms=model.terms[!grepl(x=model.terms, pattern="I(", fixed=T)]
		model.terms=model.terms[!grepl(x=model.terms, pattern="^2)", fixed=T)]
		#exclude interactions and squared terms from model.terms:
		#create new data to be used to determine fitted values:
		ii.data=model.res@frame
		all.fitted=lapply(use.list, function(use){
			if(length(circ.var.name)==1){
				set.circ.var.to.zero=sum(circ.var.name%in%use)==0
			}else{
				set.circ.var.to.zero=F
			}
			
			if(length(use)==0){use=model.terms}
			new.data=vector("list", length(model.terms))
			usel=model.terms%in%use
			#if(length(use)>0)
			for(i in 1:length(model.terms)){
				if(is.factor(ii.data[, model.terms[i]])){
					new.data[[i]]=levels(ii.data[, model.terms[i]])
				}else if(!is.factor(ii.data[, model.terms[i]]) & usel[i] & ifelse(length(circ.var.name)==0, T, !grepl(x=model.terms[i], pattern=circ.var.name))){
					new.data[[i]]=seq(from=min(ii.data[, model.terms[i]]), to=max(ii.data[, model.terms[i]]), length.out=resol)
				}else  if(!is.factor(ii.data[, model.terms[i]]) & ifelse(length(circ.var.name)==0, T, !grepl(x=model.terms[i], pattern=circ.var.name))){
					new.data[[i]]=mean(ii.data[, model.terms[i]])
				}
			}
			names(new.data)=model.terms
			if(length(circ.var.name)==1){
				new.data=new.data[!(model.terms%in%paste(c("sin(", "cos("), circ.var.name, ")", sep=""))]
				if(sum(grepl(pattern=circ.var.name, x=use))>0){
					new.data=c(new.data, list(seq(min(circ.var, na.rm=T), max(circ.var, na.rm=T), length.out=resol)))
					names(new.data)[length(new.data)]=circ.var.name
				}else{
					new.data=c(new.data, list(0))
				}
				model.terms=model.terms[!(model.terms%in%paste(c("sin(", "cos("), circ.var.name, ")", sep=""))]
			}
			xnames=names(new.data)
			#browser()
			new.data=data.frame(expand.grid(new.data))
			names(new.data)=xnames
			if(length(circ.var.name)==1){
				names(new.data)[ncol(new.data)]=circ.var.name
			}
			if(set.all.effects.2.zero){
				for(iterm in setdiff(colnames(new.data), c("(Intercept)", use))){
					new.data[, iterm]=0
				}
			}
			m.mat=model.matrix(object=as.formula(paste(c("~", model), collapse="")), data=new.data)
			if(set.circ.var.to.zero){
				m.mat[,paste(c("sin(", circ.var.name, ")"), collapse="")]=0
				m.mat[,paste(c("cos(", circ.var.name, ")"), collapse="")]=0
			}
			#get CIs for fitted values:
			ci=lapply(all.res, function(x){
				#return(apply(m.mat[names(fixef(model.res)), , drop=F]*as.vector(x[, names(fixef(model.res))]), 2, sum))
				return(m.mat[, names(fixef(model.res)), drop=F]%*%as.vector(x[, names(fixef(model.res))]))
			})
			ci=matrix(unlist(ci), ncol=nboots, byrow=F)
			ci=t(apply(ci, 1, quantile, prob=c((1-level)/2, 1-(1-level)/2), na.rm=T))
			colnames(ci)=c("lower.cl", "upper.cl")
			fv=m.mat[, names(fixef(model.res))]%*%fixef(model.res)
			if(class(model.res)[[1]]!="lmerMod"){
				if(model.res@resp$family$family=="binomial"){
					ci=exp(ci)/(1+exp(ci))
					fv=exp(fv)/(1+exp(fv))
				}else if(model.res@resp$family$family=="poisson" | substr(x=model.res@resp$family$family, start=1, stop=17)=="Negative Binomial"){
					ci=exp(ci)
					fv=exp(fv)
				}
			}
			return(data.frame(new.data, fitted=fv, ci))
		})
		result=all.fitted
		names(result)=unlist(lapply(use.list, paste, collapse="@"))
	}else{
		result=NULL
	}
	if(!is.list(use)){result=as.data.frame(result[[1]])}
	all.boots=matrix(unlist(all.res), nrow=nboots, byrow=T)
	colnames(all.boots)=colnames(all.res[[1]])
	ci.est=apply(all.boots, 2, quantile, prob=c((1-level)/2, 1-(1-level)/2), na.rm=T)
	if(length(fixef(model.res))>1){
		ci.est=data.frame(orig=fixef(model.res), t(ci.est)[1:length(fixef(model.res)), ])
	}else{
		ci.est=data.frame(orig=fixef(model.res), t(t(ci.est)[1:length(fixef(model.res)), ]))
	}
	return(list(ci.estimates=ci.est, ci.fitted=result, warnings=all.warns, ind.boots=all.boots))
}


###########################################################################################################
###########################################################################################################
boot.glmm<-function(model.res, excl.warnings=F, nboots=1000, para=F, use.u=F, n.cores=c("all-1", "all"), save.path=NULL){
	n.cores=n.cores[1]
	keepWarnings<-function(expr) {
		localWarnings <- list()
		value <- withCallingHandlers(expr,
			warning = function(w) {
				localWarnings[[length(localWarnings)+1]] <<- w
				invokeRestart("muffleWarning")
			}
		)
		list(value=value, warnings=localWarnings)
	}
	if(excl.warnings){
		boot.fun<-function(x, model.res., keepWarnings., use.u., save.path.){
			xdone=F
			while(!xdone){
				i.res=keepWarnings.(bootMer(x=model.res., FUN=fixef, nsim=1, use.u=use.u.)$t)
				if(length(unlist(i.res$warnings)$message)==0){
					xdone=T
				}
			}
			if(length(save.path.)>0){
				i.res=i.res$value
				save(file=paste(c(save.path., "/b_", x, ".RData"), collapse=""), list="i.res")
			}
			return(i.res$value)
		}
	}else{
		boot.fun<-function(x, model.res., keepWarnings., use.u., save.path.){
			i.res=bootMer(x=model.res, FUN=fixef, nsim=1, use.u=use.u.)$t
			if(length(save.path.)>0){
				save(file=paste(c(save.path., "/b_", x, ".RData"), collapse=""), list="i.res")
			}
			return(i.res)
		}
	}
	if(para){
		require(parallel)
    cl <- makeCluster(getOption("cl.cores", detectCores()))
		if(n.cores!="all"){
			if(n.cores!="all-1"){n.cores=length(cl)-1}
			if(n.cores<length(cl)){
				cl=cl[1:n.cores]
			}
		}
    parLapply(cl=cl, 1:length(cl), fun=function(x){
      library(lme4)
      return(invisible(""))
    })
    all.coeffs=parLapply(cl=cl, X=1:nboots, fun=boot.fun, model.res.=model.res, keepWarnings.=keepWarnings, use.u=use.u, save.path.=save.path)
    parLapply(cl=cl, X=1:length(cl), fun=function(x){rm(list=ls())})
    stopCluster(cl)
	}else{
    all.coeffs=lapply(X=1:nboots, FUN=boot.fun, model.res.=model.res, keepWarnings.=keepWarnings, use.u=use.u, save.path.=save.path)
	}
	ci=matrix(unlist(all.coeffs), nrow=length(all.coeffs), byrow=T)
	colnames(ci)=names(fixef(model.res))
	ci=apply(ci, 2, quantile, prob=c(0.025, 0.975), na.rm=T)
	ci=data.frame(orig=fixef(model.res), t(ci))
	return(ci)
}
```
