## Supplementary material for "Male Guinea baboons are oblivious to their females’ whereabouts": Data Analyses: coxme_stab.r

```
coxme.stab<-function(mres, data, formula, para=F, n.cores=c("all-1", "all"), contr=NULL){
	if(is.null(contr)){contr=coxme.control()}
	ranef2=unique(names(mres$vcoef))
	to.do=lapply(ranef2, function(x){sort(unique(as.character(data[, x])))})
	to.do=data.frame(
		ranef=rep(ranef2, times=unlist(lapply(to.do, length))),
		level=unlist(to.do)
	)
	to.do$ranef=as.character(to.do$ranef)
	to.do$level=as.character(to.do$level)
	fit.fun<-function(i, to.do., data., formula., mres.=mres){
		idata=subset(data, data.[, to.do.$ranef[i]]!=to.do.$level[i])
		im=try(coxme(as.formula(formula.), data=idata, control=contr), silent=T)
		if(class(im)[[1]]!="try-error"){
			return(c(nrow(idata), coefficients(im), unlist(im$vcoef)))
		}else{
			return(c(nrow(idata), rep(NA, length(coefficients(mres.))), rep(NA, length(unlist(mres.$vcoef)))))
		}
	}
	if(!para){
		xx=lapply(X=1:nrow(to.do), FUN=fit.fun, to.do.=to.do, data.=data, formula.=formula)
	}else{
    library(parallel)
    cl <- makeCluster(getOption("cl.cores", detectCores()))
    n.cores=n.cores[1]
		if(n.cores!="all"){
			if(n.cores!="all-1"){n.cores=length(cl)-1}
			if(n.cores<length(cl)){
				cl=cl[1:n.cores]
			}
		}
    parLapply(cl=cl, 1:length(cl), fun=function(x){
      library(coxme)
      return(invisible(""))
    })
		xx=parLapply(X=1:nrow(to.do), cl=cl, fun=fit.fun, to.do.=to.do, data.=data, formula.=formula)
    parLapply(cl=cl, X=1:length(cl), fun=function(x){rm(list=ls())})
    stopCluster(cl)
	}
	xx=matrix(unlist(xx), nrow=length(xx), byrow=T)
	colnames(xx)=c("N", names(coefficients(mres)), paste(names(mres$vcoef), unlist(lapply(mres$vcoef, names)), sep="@"))
	to.do=data.frame(to.do, xx)
	xsum=cbind(orig=c(coefficients(mres), unlist(mres$vcoef)), t(apply(xx[, -1], 2, range, na.rm=T)))
	colnames(xsum)[2:3]=c("min", "max")
	return(list(summary=xsum, detailed=to.do))
}
```
