## Supplementary material for "Male Guinea baboons are oblivious to their females’ whereabouts": Data Analyses: diagnostic_fcns.r

```
##functions providng various diagnostics for evaluating model assumptions and stability and
#for helping with setting an appropriate random slopes structure for (G)LMMs
#written by Roger Mundry, last modified early 2016
ranef.diagn.plot<-function(model.res, QQ=F, col=grey(0.5, alpha=0.75)){
  old.par = par(no.readonly = TRUE)
	if(class(model.res)[[1]]!="glmmTMB"){
		n.plots=sum(unlist(lapply(ranef(model.res), length)))
	}else{
		n.plots=sum(unlist(lapply(ranef(model.res)[["cond"]], ncol)))+sum(unlist(lapply(ranef(model.res)[["zi"]], ncol)))
	}
	x=ifelse(n.plots%%2==1,n.plots+1,n.plots)
	xmat=outer(1:x, 1:(1+x/2), "*")-n.plots
	colnames(xmat)=1:(1+x/2)
	rownames(xmat)=1:x
	xmat=as.data.frame(as.table(xmat))
	xmat=subset(xmat, as.numeric(as.character(xmat$Var1))>=as.numeric(as.character(xmat$Var2)) & xmat$Freq>=0)
	sum.diff=as.numeric(as.character(xmat$Var1))-as.numeric(as.character(xmat$Var2))+xmat$Freq
	xmat=xmat[sum.diff==min(sum.diff),]
	xmat=xmat[which.min(xmat$Freq),]
	par(mfrow=c(xmat$Var2, xmat$Var1))
	par(mar=c(rep(2, 3), 1))
	par(mgp=c(1, 0.5, 0))
	xnames=lapply(ranef(model.res), names)
	xnames=lapply(xnames, gsub, pattern="(Intercept)", replacement="icpt", fixed=T)
	if(class(model.res)[[1]]!="glmmTMB"){
		for(i in 1:length(ranef(model.res))){
			to.plot=ranef(model.res)[[i]]
			for(k in 1:ncol(to.plot)){
				if(QQ){
					qqnorm(to.plot[,k], main="", tcl=-0.25, xlab="", ylab="", pch=19, col=col)
					qqline(to.plot[,k])
				}else{
					hist(to.plot[,k], main="", tcl=-0.25, xlab="", ylab="", col=NA)
				}
				mtext(text=paste(c(names(ranef(model.res)[i]), xnames[[i]][k]), collapse="_"), side=3)
			}
		}
	}else{
		for(i in c("cond", "zi")){
			if(length(ranef(model.res)[[i]])>0){
				for(j in 1:length(ranef(model.res)[[i]])){
					to.plot=ranef(model.res)[[i]][[j]]
					for(k in 1:ncol(to.plot)){
						if(QQ){
							qqnorm(to.plot[,k], main="", tcl=-0.25, xlab="", ylab="", pch=19, col=col)
							qqline(to.plot[,k])
						}else{
							hist(to.plot[,k], main="", tcl=-0.25, xlab="", ylab="", col=NA)
						}
						mtext(text=paste(c(names(ranef(model.res)[i]), names(ranef(model.res)[[i]][j]), colnames(to.plot)[k]), collapse=", "), side=3)
					}
				}
			}
		}
	}
	par(old.par)
}

diagnostics.plot<-function(mod.res, col=grey(level=0.25, alpha=0.5), size.fac=1, weights=NULL){
	if(is.null(weights)){
		if(class(mod.res)[[1]]=="lm" | class(mod.res)[[1]]=="glm"){
			weights=mod.res$model$"(weights)"
			if(is.null(weights)){weights=rep(1, length(residuals(mod.res)))}
		}else if(class(mod.res)[[1]]=="lmerMod"){
			weights=mod.res@resp$weights
		}else{
			weights=rep(1, length(residuals(mod.res)))
		}
	}
	weights=size.fac*weights/max(weights)
  old.par = par(no.readonly = TRUE)
  par(mfrow=c(2, 2))
  par(mar=c(3, 3, 1, 0.5))
  hist(residuals(mod.res), probability=T, xlab="", ylab="", main="", col=NA)
  mtext(text="histogram of residuals", side=3, line=0)
  x=seq(min(residuals(mod.res)), max(residuals(mod.res)), length.out=100)
  lines(x, dnorm(x, mean=0, sd=sd(residuals(mod.res))))
  qqnorm(residuals(mod.res), main="", pch=19, col=col, cex=sqrt(weights))
  qqline(residuals(mod.res))
  mtext(text="qq-plot of residuals", side=3, line=0)
  plot(fitted(mod.res), residuals(mod.res), pch=19, col=col, cex=sqrt(weights))
  abline(h=0, lty=2)
  mtext(text="residuals against fitted values", side=3, line=0)
  par(old.par)
}

lev.thresh<-function(model.res){
	k=length(coefficients(model.res))
	n=length(residuals(model.res))
 return(2*(k+1)/n)
}

overdisp.test<-function(x){
	##last updated Nov 26 2019
	##additions/changes: bugfix for zeroinfl; deals with glmmTMB (beta and nbinom family)
	##function needed for Gamma family:
	gamma.par.transf<-function(mean=NULL, var=NULL, shape=NULL, scale=NULL){
		##parameterizations are
			##shape and scale
			##shape and mean
			##var and mean
		if(is.null(mean) & is.null(var) & !is.null(shape) & !is.null(scale)){
			mean=shape*scale
			var=shape*scale^2
		}else if(is.null(shape) & is.null(scale) & !is.null(mean) & !is.null(var)){
			scale=var/mean
			shape=mean/scale
		}else if(is.null(scale) & is.null(var) & !is.null(shape) & !is.null(mean)){
			scale=mean/shape
			var=shape*scale^2
		}else{
			stop("unsupported parameter combination")
		}
		return(list(shape=as.vector(shape), scale=as.vector(scale),  mean=as.vector(mean), var=as.vector(var)))
	}
	
  if(class(x)[[1]]!="glmmTMB"){
		pr=residuals(x, type ="pearson")
		sum.dp=sum(pr^2)
	}
  n.model.terms=NULL
  if(class(x)[[1]]=="glmerMod"){
    n.model.terms=length(fixef(x))+nrow(as.data.frame(summary(x)$varcor))
    if(grepl(x=as.character(x@call)[4], pattern="negative.binomial") | grepl(x=as.character(x@call)[4], pattern="Gamma")){n.model.terms=n.model.terms+1}
	}else if(class(x)[[1]]=="glm" | class(x)[[1]]=="negbin"){
    n.model.terms=length(x$coefficients)
    n.model.terms=n.model.terms+length(summary(x)$theta)
  }else if(class(x)[[1]]=="zeroinfl"){
		n.model.terms=sum(unlist(lapply(x$coefficients, length)))
		n.model.terms=n.model.terms+length(summary(x)$theta)
  }else if(class(x)[[1]]=="betareg"){
		n.model.terms=sum(unlist(lapply(x$coefficients, length)))
  }else if(class(x)[[1]]=="glmmTMB"){
		print("caution: support for models of class glmmTMB is experimental and embryonic")
		print("and don't worry if it takes some time")
		#browser()
		#n.model.terms=sum(unlist(lapply(fixef(x), length)))+nrow(extract.ranef.from.glmmTB(x))
		n.model.terms=sum(unlist(lapply(fixef(x), length)))##fixed effects
		n.model.terms=n.model.terms+sum(unlist(lapply(summary(x)$varcor, function(y){
			if(!is.null(y)){
				lapply(y, function(yy){
					xx=dim(attr(yy, "correlation"))
					return((xx[1]^2-xx[1])/2+length(attr(yy, "stddev")))
				})
			}else{
				0
			}
		})))
		if((grepl(x=x$modelInfo$family$family, pattern="nbinom") | x$modelInfo$family$family=="beta") &
			length(attr(terms(x$call$dispformula), "term.labels"))==0){
			#n.model.terms=n.model.terms+1
			##but maybe it needs to be done for negbin??? need to check this
		}
		pr=residuals(x, type ="response")
		if(x$modelInfo$family$family=="poisson"){
			fitted.var = fitted(x)
		}else{
			#model.terms=attr(terms(x$call$dispformula), "term.labels")#attr(terms(x$call$dispformula), "intercept")
			sigma.f=exp(as.vector(model.matrix(object=x$call$dispformula, data=x$frame)[, names(fixef(x)$disp), drop=F]%*%fixef(x)$disp))
			if(x$modelInfo$family$family=="nbinom2"){
				fitted.var = fitted(x)*(1+fitted(x)/sigma.f)#fitted(x) + fitted(x)^2/sigma(x)
			}else if(x$modelInfo$family$family=="nbinom1"){
				fitted.var = fitted(x)*(1+sigma.f)#fitted(x) + fitted(x)/sigma(x)
			}else if(x$modelInfo$family$family=="beta"){
				xfitted=fitted(x)
				shape1 = xfitted * sigma.f
				shape2 = (1 - xfitted) * sigma.f
				#xfitted(1-xfitted)/(1+sigma(x))#according to the glmmTMB help page for sigma
				fitted.var = shape1*shape2/((shape1+shape2)^2*(shape1+shape2+1))
			}else if(x$modelInfo$family$family=="Gamma"){
				#x.disp=exp(as.vector(model.matrix(object=x$call$dispformula, data=x$frame)%*%log(sigma(x))))
				xx=gamma.par.transf(mean=fitted(x), var=NULL, shape=1/sigma.f, scale=NULL)
				fitted.var = xx$var
				#xdf=length(residuals(x))-(length(coef(x))+1)
				#n.model.terms=sum(unlist(lapply(fixef(x), length)))+sum(unlist(lapply(summary(x)$varcor, function(y){length(as.data.frame(y))})))
				xdf=length(residuals(x))-n.model.terms
			}
		}
		#n.model.terms=sum(unlist(lapply(fixef(x), length)))+sum(unlist(lapply(summary(x)$varcor, function(y){length(as.data.frame(y))})))
		#xdf=length(pr)-n.model.terms
		sum.dp=sum((pr/sqrt(fitted.var))^2)
		#if(as.numeric(substr(sessionInfo()[["otherPkgs"]][["glmmTMB"]]$Version, start=1, stop=1))>0){
			#sum.dp=sum(residuals(x, type ="pearson")^2)
		#}
  }
  if(!is.null(n.model.terms)){
		xdf=length(pr)-n.model.terms
		return(data.frame(chisq=sum.dp, df=xdf, P=1-pchisq(sum.dp, xdf), dispersion.parameter=sum.dp/xdf))
	}else{
		print("model isn't of any of the currently supported classes (glm, negbin, zeroinfl, glmerMod, glmmTMB)")
	}
}

fe.re.tab<-function(fe.model, re, other.vars=NULL, data, treat.covs.as.factors=c(NA, F, T)){
	print("please read the documentation in the beginning of the script")
	#function helping in determinining which random slopes are needed
	#last updated: 2018, June 6
	#latest updates:
		#major revision of how interactions are treated; reveals in the summary
			#for combinations factors whether the number of observations is >1 (seperately for each level of the random effect)
			#for combination of covariates whether the number of unique combinations per level of random effect is >2
			#for combinations of factors and covariates whether the number unique values per covariate is larger than 2 (when treated as covariate)
				#or larger than 1 (when treated as a factor), separately for each commbination of levels of fixed and random effects factors
		#treat.covs.as.factors can be NA in which case covariates aretreates as such and as factors
			#this is the default now
		#output gives information about which fixed effects are covariates and factors aso for interactions
			#and also how covariates were treated
	#input/arguments:
		#data: a dataframe with all relevant variables (including the response)
		#fe.model: character; the model wrt the fixed effect (including the response); e.g., "r~f1*c*f2"
		#re: character; either a vector with the names of the random effects (e.g., c("re1", "re2")) or a random intercepts expression (e.g., "(1|re1)+(1|re2)")
		#other.vars: character, optional; a vector with the names of variables which are to be kept in the data considered and returned
		#treat.covs.as.factors: logical, when set to TRUE covariates will be treated like factors (see value/summary for details)
	#value: list with the following entries:
		#detailed: list with cross-tabulations ffor each combination of (main) fixed and random effect 
		#summary: list tables...
			#telling for each combination of (main) fixed and random effect...
				#the number of levels of the random effect with a given number of unique values of the fixed effect (in case of a covariate)
				#the number of levels of the random effect with a given number of levels of the fixed effect for which at least two cases do exist (in case of a factor)
			#telling for each combination of interaction and random effect...
				#the combination of the above two informations, i.e., the number of individuals with a given number of unique values of the covariate
					#and a given number of factor levels for which at least two cases exist
		#data: data frame containing all relevant variables (i.e., response, fixed and random effects as well as those indicated in other.vars (e.g., offset terms)
			#also includes columns for dummy variables coding the levels (except the reference level) of all factors
		#pot.terms: length one vector comprising the model wrt the random slopes (for all combinations of fixed and random effects, i.e., most likely some will need to be omitted)
			#note that this comprises only the random slopes but not the correlations between random slopes and intercepts not the random intercept itself
		#pot.terms.with.corr: length one vector comprising the model wrt the random slopes (for all combinations of fixed and random effects, i.e., most likely some will need to be omitted)
			#note that this comprises the random slopes and intercepts  and also all correlations among them
	treat.covs.as.factors=treat.covs.as.factors[1]
	if(sum(grepl(x=re, pattern="", fixed=T))>0 & length(re)==1){#if random effects are handed over as formula
		re=gsub(x=re, pattern="(1|", replacement="", fixed=T)
		re=gsub(x=re, pattern=")", replacement="", fixed=T)
		re=gsub(x=re, pattern=" ", replacement="", fixed=T)
		re=unlist(strsplit(re, split="+", fixed=T))
	}
	fe.model=gsub(x=fe.model, pattern=" ", replacement="", fixed=T)#remove spaces
	model.terms=attr(terms(as.formula(fe.model)), "term.labels")#get individual terms from fixed effects model
	fe.me=model.terms[!grepl(x=model.terms, pattern=":", fixed=T)]#remove interactions
	fe.me=fe.me[!grepl(x=fe.me, pattern="^", fixed=T)]#remove squares terms
	resp=unlist(strsplit(fe.model, split="~", fixed=T))[1]#determine response
	if(substr(resp, start=1, stop=6)=="cbind("){
		resp=gsub(x=resp, pattern="cbind(", replacement="", fixed=T)
		resp=gsub(x=resp, pattern=")", replacement="", fixed=T)
		resp=gsub(x=resp, pattern=" ", replacement="", fixed=T)
		resp=unlist(strsplit(resp, split=",", fixed=T))
	}
	xx=setdiff(c(resp, fe.me, re, other.vars), names(data))
	if(length(xx)>0){
		stop(paste(c("error: preditor(s) missing in the data is/are ", paste(xx, collapse=", ")), collapse=""))
	}
	data=droplevels(as.data.frame(na.omit(data[, c(resp, fe.me, re, other.vars)])))#keep complete data wrt all relevant variables
	model.terms=model.terms[!grepl(x=model.terms, pattern="^", fixed=T)]#remove squares terms
	modes=rep(NA, length(model.terms))#initialize vector storing whether PVs are factors or not
	effect=c(rep("main", length(fe.me)), rep("int", length(model.terms)-length(fe.me)))#create vector telling for each model term whether it is a main effect of not
	for(i in 1:ncol(data)){#for all columns in data
		if(is.character(data[, i])){data[, i]=as.factor(data[, i])}#turn character column into factor
		modes[i]=class(data[, i])#and determine its class
	}
	names(modes)=names(data)#name 'm
	to.do=data.frame(expand.grid(re=re, fe=fe.me))#create data frame with one column for each combination of fixed main and random effect
	to.do$re=as.character(to.do$re)#reformat to character (for later addressing by name)
	to.do$fe=as.character(to.do$fe)#reformat to character (for later addressing by name)
	res.detailed=lapply(1:nrow(to.do), function(xrow){#create detailed results by lapply-ing over the rows of to.do
		table(data[,to.do$re[xrow]], data[,to.do$fe[xrow]])#cross tabulate the respective fixed and random effect
	})
	names(res.detailed)=paste(to.do$fe, to.do$re, sep="_within_")#name it
	modes=modes[as.character(to.do$fe)]
	modes=gsub(x=modes, pattern="integer", replacement="numeric")
	to.do$modes.cons=modes
	to.do$modes=modes
	if(is.na(treat.covs.as.factors) & any(to.do[, "modes.cons"]=="numeric")){
		xx=to.do[to.do[, "modes.cons"]=="numeric", ]
		xx["modes.cons"]="factor"
		to.do=rbind(to.do, xx)
		to.do=to.do[order(to.do[, "re"], to.do[, "fe"]), ]
	}else if(!is.na(treat.covs.as.factors) & treat.covs.as.factors){
		to.do$modes.cons="factor"
	}
	to.do=to.do[order(to.do$fe), ]
	res.summary=lapply(1:nrow(to.do), function(xrow){#begin with creating summary results by lapply-ing over the rows of to.do
		#browser()
		xwhat=paste(to.do[xrow, c("fe", "re")], collapse="_within_")
		if(to.do$modes.cons[xrow]!="factor"){#if fixed effect is not a factor
			ires=table(apply(res.detailed[[xwhat]]>0, 1, sum))#determine number of levels of the random effect per number of unique cases of the fixed effect
		}else{#if fixed effect is a factor
			ires=table(apply(res.detailed[[xwhat]]>1, 1, sum))#determine number of levels of the random effect per number of unique cases of the fixed effect with at least two observations
		}
		ires=c(ires, tot=nrow(res.detailed[[xwhat]]))
	})
	xx=to.do[, "modes.cons"]
	xx[xx=="numeric"]="covariate"
	xx[xx=="integer"]="covariate"
	xx[to.do[, "modes.cons"]!=to.do[, "modes"]]="covariate as factor"
	xx=paste("(", xx, ")", sep="")
	names(res.summary)=paste(paste(to.do$fe, to.do$re, sep="_within_"), xx, sep=" ")#name it
	#append 'factor' or 'covariate' to the names:
	#names(res.summary)[modes[to.do$fe]=="factor"]=paste(names(res.summary)[modes[to.do$fe]=="factor"], "(factor)", sep=" ")
	#names(res.summary)[!modes[to.do$fe]=="factor"]=paste(names(res.summary)[!modes[to.do$fe]=="factor"], "(covariate)", sep=" ")
	to.do=data.frame(expand.grid(re=re, int=setdiff(model.terms, fe.me)))#create data frame with one row for each combination of interaction and random effect
	if(nrow(to.do)>0){
		#add two columns denoting original mode:
		to.do$mode=unlist(lapply(strsplit(as.character(to.do$int), split=":", fixed=T), function(x){
			paste(modes[x], collapse=":")
		}))
		to.do$mode.cons=to.do$mode
		if(is.na(treat.covs.as.factors)){#if is.na(treat.covs.as.factors)
			xx=to.do[grepl(x=to.do$mode, pattern="numeric"), ]#add rows duplicating interactions involving covariates (to treat them as covariate and factor
			xx$mode.cons=gsub(x=xx$mode.cons, pattern="numeric", replacement="factor")
			to.do=rbind(to.do, xx)
			to.do=to.do[order(to.do$re, to.do$int), ]
		}else if(treat.covs.as.factors){#if treat.covs.as.factors
			to.do$mode.cons=gsub(x=to.do$mode.cons, pattern="numeric", replacement="factor")#change mode to be considered
		}
		to.do[, "int"]=as.character(to.do[, "int"])#reformat to character (for later addressing by name)
		to.do[, "re"]=as.character(to.do[, "re"])#reformat to character (for later addressing by name)
		##this needs to be rewritten!!!!!!!!!!!!!!!!!!!!!!!!!!!!!!
		to.add=lapply(1:nrow(to.do), function(xrow){#treat combinations of fixed and random effect by lapply-ing over the rows of to.do
			#browser()
			iterms=unlist(strsplit(to.do[xrow, "int"], split=":", fixed=T))#determine main effects involved in the interaction...
			imodes=unlist(strsplit(to.do[xrow, "mode.cons"], split=":", fixed=T))#... and their classes
			imodes.orig=unlist(strsplit(to.do[xrow, "mode"], split=":", fixed=T))#... and their classes
			if(all(imodes.orig=="factor")){
				comb.fac=lapply(iterms[imodes=="factor"], function(x){paste(x, data[, x], sep="")})
				comb.fac=matrix(unlist(comb.fac), ncol=length(comb.fac), byrow=F)
				comb.fac=apply(comb.fac, 1, paste, collapse="__")
				ires=table(data[,to.do$re[xrow]], comb.fac)
				class(ires)="matrix"
				ires.summary=aggregate(1:nrow(ires), data.frame(ires>1), length)
				xx=as.matrix(ires.summary[, -ncol(ires.summary)])
				xx[xx]=">1"
				xx[xx!=">1"]="!>1"
				ires.summary=data.frame(xx, freq=ires.summary$x)
				ires=list(detailed=ires, summary=ires.summary)
			}else if(any(imodes.orig=="factor")){
				comb.fac=lapply(iterms[imodes.orig=="factor"], function(x){paste(x, data[, x], sep="")})
				comb.fac=matrix(unlist(comb.fac), ncol=length(comb.fac), byrow=F)
				comb.fac=apply(comb.fac, 1, paste, collapse="__")
				covs=lapply(iterms[imodes.orig=="numeric"], function(x){data[, x]})
				if(all(imodes=="factor")){
					ires=lapply(covs, function(x){
						x=tapply(x, list(data[, to.do[xrow, "re"]], comb.fac), function(x){
							sum(table(x)>1)
						})
						x[is.na(x)]=0
						return(x)
					})
				}else{
					ires=lapply(covs, function(x){
						x=tapply(x, list(data[, to.do[xrow, "re"]], comb.fac), function(x){
							length(unique(x))
						})
						x[is.na(x)]=0
						return(x)
					})
				}
				xx=matrix(unlist(lapply(ires, c)), ncol=length(ires), byrow=F)
				xx=apply(xx, 1, paste, collapse="/")
				xx=matrix(xx, ncol=ncol(ires[[1]]), byrow=F)
				colnames(xx)=colnames(ires[[1]])
				rownames(xx)=rownames(ires[[1]])
				ires.detailed=xx
				if(all(imodes=="factor")){
					xx=matrix(unlist(lapply(ires, c))>=2, ncol=length(ires), byrow=F)
					xx[xx]=">1"
					xx[xx!=">1"]="!>1"
				}else{
					xx=matrix(unlist(lapply(ires, c))>=3, ncol=length(ires), byrow=F)
					xx[xx]=">2"
					xx[xx!=">2"]="!>2"
				}
				xx=apply(xx, 1, paste, collapse="/")
				xx=matrix(xx, ncol=ncol(ires[[1]]), byrow=F)
				colnames(xx)=colnames(ires[[1]])
				rownames(xx)=rownames(ires[[1]])
				ires.summary=aggregate(1:nrow(xx), data.frame(xx), length)
				colnames(ires.summary)[ncol(ires.summary)]="freq"
				ires=list(detailed=ires.detailed, summary=ires.summary)
			}else{
				xx=lapply(iterms, function(x){
					apply(res.detailed[[paste(c(x, to.do[xrow, "re"]), collapse="_within_")]]>0, 1, sum)
				})
				yy=matrix(unlist(xx), ncol=length(xx), byrow=F)
				colnames(yy)=iterms
				rownames(yy)=names(xx[[1]])
				ires=yy
				ires.summary=aggregate(1:nrow(ires), data.frame(ires), length)
				colnames(ires.summary)=c(colnames(ires), "freq")#[ncol(ires.summary)]
				ires=list(detailed=ires, summary=ires.summary)
				#browser()
			}
			return(ires)
		})
		xx=paste(to.do$int, to.do$re, sep="_within_")#gsub(x=paste(to.do$int, to.do$re, sep="_within_"), pattern=":", replacement="_", fixed=T)#name it
		mode.mat=to.do[, "mode"]
		mode.mat=gsub(x=mode.mat, pattern="numeric", replacement="COV")
		mode.mat=gsub(x=mode.mat, pattern="factor", replacement="FAC")
		mode.mat=strsplit(mode.mat, split=":", fixed=T)#), nrow=nrow(to.do), byrow=T)
		mode.cons.mat=to.do[, "mode.cons"]
		mode.cons.mat=gsub(x=mode.cons.mat, pattern="numeric", replacement="COV")
		mode.cons.mat=gsub(x=mode.cons.mat, pattern="factor", replacement="FAC")
		mode.cons.mat=strsplit(mode.cons.mat, split=":", fixed=T)#), nrow=nrow(to.do), byrow=T)
		for(i in 1:length(mode.mat)){
			mode.mat[[i]][mode.mat[[i]]!=mode.cons.mat[[i]]]="COVasFAC"
		}
		#mode.mat[mode.mat=="numeric"]="COV"
		#mode.mat[mode.mat=="factor"]="FAC"
		#mode.mat[mode.mat!=mode.cons.mat]="COVasFAC"
		xx=paste(xx, paste("(", unlist(lapply(mode.mat, paste, collapse=":")), ")", sep=""), sep=" ")
		to.add.2=lapply(to.add, "[[", "summary")
		names(to.add.2)=xx
		res.summary=c(res.summary, to.add.2)#and append to.add to res.summary
		to.add.2=lapply(to.add, "[[", "detailed")
		names(to.add.2)=xx
		res.detailed=c(res.detailed, to.add.2)
	}
	#add columns with the dummy coded factor levels to data:
	to.code=fe.me[modes[fe.me]=="factor"]#determine factors among the fixed effects:
	if(length(to.code)>0){
		coded=lapply(to.code, function(xc){#for each factor
			lapply(levels(data[, xc])[-1], function(xl){#for all levels except the reference level
				as.numeric(data[, xc]==xl)#code it
			})
		})
		coded=matrix(unlist(coded), nrow=nrow(data), byrow=F)#reformat to matrix
		xnames=unlist(lapply(to.code, function(xc){#determine column names to be given to the matrix...
			paste(xc, levels(data[, xc])[-1], sep=".")
		}))
		colnames(coded)=xnames#... and use 'm
		data=data.frame(data, coded)#and append 'm to data
	}else{
		xnames=""
	}
	#now the model expression wrt random slopes; first no correllations between random slopes and intercepts:
	pot.terms=c(xnames, fe.me[modes[fe.me]!="factor"])#create vector with names of dummy variables and names of fixed main effects not being factors
	pot.terms=outer(pot.terms, re, Vectorize(function(x, y){#create matrix with separate model term for each combination of fixed and random effect
		paste(c("(0+", x, "|", y, ")"), collapse="")
	}))
	pot.terms=paste(c(t(pot.terms)), collapse="+")#put them all in a single entry
	#now the random slopes part of the model including random intercepts and slopes and their correlation:
	pot.terms.with.corr=paste(c(xnames, fe.me[modes[fe.me]!="factor"]), collapse="+")#get all fixed effects terms together...
	pot.terms.with.corr=paste(paste("(1+", pot.terms.with.corr, "|", re, ")", sep=""), collapse="+")#... and paste random effects, brackets and all that 
		#(and everything in a single entry)
	return(list(detailed=res.detailed, summary=res.summary, data=data, pot.terms=pot.terms, pot.terms.with.corr=pot.terms.with.corr))
}

write.fe.re.summary2file<-function(x, file){
	c.tab<-function(x, incl.fst=F, incl.rownames=T){
		res=format(as.matrix(x))
		res=gsub(x=res, pattern=" ", replacement="", fixed=T)
		res=rbind(colnames(res), res)
		rownames(res)=NULL
		colnames(res)=NULL
		res[is.na(res)]="NA"
		n.char.range=apply(res, 2, function(x){range(nchar(x))})
		#res[, 1]=unlist(lapply(res[, 1], function(x){paste(c(x, rep(" ", n.char.range[2, 1]-nchar(x))), collapse="")}))
		res=matrix(unlist(lapply(1:ncol(res), function(x){
			unlist(lapply(res[, x], function(y){paste(c(rep(" ", n.char.range[2, x]-nchar(y)), y), collapse="")}))
		})), nrow=nrow(res), byrow=F)
		res=apply(res, 1, paste, collapse=paste(rep(" ", 1), collapse=""))
		return(res)
	}

	append=F
	for(i in 1:length(x)){
		write.table(x=names(x[i]), file=file, row.names=F, col.names=F, sep="\t", quote=F, append=append)
		append=T
		xx=x[[i]]
		if(is.null(dim(xx))){
			xx=cbind(names(xx), xx)
			colnames(xx)=NULL
			xx=c.tab(xx)
		}else{
			rownames(xx)=NULL
			xx=c.tab(x=xx)
		}
		xx=matrix(xx, ncol=1)
		write.table(x=xx, file=file, row.names=F, col.names=F, sep="\t", quote=F, append=append)
		write.table(x="", file=file, row.names=F, col.names=F, sep="\t", quote=F, append=append)
	}
}

m.stab.plot<-function(est, lower=NULL, upper=NULL, xnames=NULL, col="black", center.at.null=F, reset.par=T, pch=18, xlim=NULL){
	#version from Aug 12 2018
	#function to plot model stability or CIs (not really supposed to be nice, but to give a quick overview/rapid diagnostic)
	#input:
		#either a three columns data frame or matrix (with rownames) handed over to argument est (first column needs to comprise the original estimate)
		#or several vectors handed over as follows:
			#est: numeric; estimated coefficients of the model
			#lower: numeric; lower limits of the estimates (either from model stability of from bootstrap)
			#upper: numeric; upper limits of the estimates (either from model stability of from bootstrap)
			#xnames: character; names to be depicted besides the error bars
		#col: character; name of the color with which data should be depicted
	if(ncol(est)==3){
		lower=est[, 2]
		upper=est[, 3]
		xnames=rownames(est)
		est=est[, 1]
		x.at=est
	}
	if(is.null(xlim)){
		xlim=range(c(0, lower, upper))
	}
  old.par = par(no.readonly = TRUE)
	par(mar=c(3, 0.5, 0.5, 0.5), mgp=c(1, 0.4, 0), tcl=-0.2)
	plot(x=est, y=1:length(est), pch=pch, xlab="estimate", ylab="", yaxt="n", xlim=xlim, type="n", ylim=c(1, length(est)+1))
	abline(v=0, lty=3)
	if(center.at.null){
		x.at=rep(0, length(est))
		text(labels=xnames, x=x.at, y=(1:length(est))+0.3, cex=0.8)
	}else{
		xx=range(est)#c(lower, upper))
		for(i in 1:length(est)){
			text(labels=xnames[i], x=x.at[i], y=i+0.3, cex=0.8, adj=c((est[i]-min(xx))/diff(xx), 0))#pos=c(2, 4)[1+as.numeric(est<0)])
		}
	}
	points(x=est, y=1:length(est), pch=pch, col=col)
	segments(x0=lower, x1=upper, y0=1:length(est), y1=1:length(est), col=col)
	if(reset.par){par(old.par)}
}

fe.re.tab.old<-function(fe.model, re, other.vars=NULL, data, treat.covs.as.factors=F){
	#function helping in determinining which random slopes are needed
	#last updated: 2015, Nov 25
	#input/arguments:
		#data: a dataframe with all relevant variables (including the response)
		#fe.model: character; the model wrt the fixed effect (including the response); e.g., "r~f1*c*f2"
		#re: character; either a vector with the names of the random effects (e.g., c("re1", "re2")) or a random intercepts expression (e.g., "(1|re1)+(1|re2)")
		#other.vars: character, optional; a vector with the names of variables which are to be kept in the data considered and returned
		#treat.covs.as.factors: logical, when set to TRUE covariates will be treated like factors (see value/summary for details)
	#value: list with the following entries:
		#detailed: list with cross-tabulations ffor each combination of (main) fixed and random effect 
		#summary: list tables...
			#telling for each combination of (main) fixed and random effect...
				#the number of levels of the random effect with a given number of unique values of the fixed effect (in case of a covariate)
				#the number of levels of the random effect with a given number of levels of the fixed effect for which at least two cases do exist (in case of a factor)
			#telling for each combination of interaction and random effect...
				#the combination of the above two informations, i.e., the number of individuals with a given number of unique values of the covariate
					#and a given number of factor levels for which at least two cases exist
		#data: data frame containing all relevant variables (i.e., response, fixed and random effects as well as those indicated in other.vars (e.g., offset terms)
			#also includes columns for dummy variables coding the levels (except the reference level) of all factors
		#pot.terms: length one vector comprising the model wrt the random slopes (for all combinations of fixed and random effects, i.e., most likely some will need to be omitted)
			#note that this comprises only the random slopes but not the correlations between random slopes and intercepts not the random intercept itself
		#pot.terms.with.corr: length one vector comprising the model wrt the random slopes (for all combinations of fixed and random effects, i.e., most likely some will need to be omitted)
			#note that this comprises the random slopes and intercepts  and also all correlations among them
	if(sum(grepl(x=re, pattern="", fixed=T))>0 & length(re)==1){#if random effects are handed over as formula
		re=gsub(x=re, pattern="(1|", replacement="", fixed=T)
		re=gsub(x=re, pattern=")", replacement="", fixed=T)
		re=gsub(x=re, pattern=" ", replacement="", fixed=T)
		re=unlist(strsplit(re, split="+", fixed=T))
	}
	fe.model=gsub(x=fe.model, pattern=" ", replacement="", fixed=T)#remove spaces
	model.terms=attr(terms(as.formula(fe.model)), "term.labels")#get individual terms from fixed effects model
	fe.me=model.terms[!grepl(x=model.terms, pattern=":", fixed=T)]#remove interactions
	fe.me=fe.me[!grepl(x=fe.me, pattern="^", fixed=T)]#remove squares terms
	resp=unlist(strsplit(fe.model, split="~", fixed=T))[1]#determine response
	if(substr(resp, start=1, stop=6)=="cbind("){
		resp=gsub(x=resp, pattern="cbind(", replacement="", fixed=T)
		resp=gsub(x=resp, pattern=")", replacement="", fixed=T)
		resp=gsub(x=resp, pattern=" ", replacement="", fixed=T)
		resp=unlist(strsplit(resp, split=",", fixed=T))
	}
	xx=setdiff(c(resp, fe.me, re, other.vars), names(data))
	if(length(xx)>0){
		stop(paste(c("error: preditor(s) missing in the data is/are ", paste(xx, collapse=", ")), collapse=""))
	}
	data=droplevels(as.data.frame(na.omit(data[, c(resp, fe.me, re, other.vars)])))#keep complete data wrt all relevant variables
	model.terms=model.terms[!grepl(x=model.terms, pattern="^", fixed=T)]#remove squares terms
	modes=rep(NA, length(model.terms))#initialize vector storing whether PVs are factors or not
	effect=c(rep("main", length(fe.me)), rep("int", length(model.terms)-length(fe.me)))#create vector telling for each model term whether it is a main effect of not
	for(i in 1:ncol(data)){#for all columns in data
		if(is.character(data[, i])){data[, i]=as.factor(data[, i])}#turn character column into factor
		modes[i]=class(data[, i])#and determine its class
	}
	names(modes)=names(data)#name 'm
	to.do=data.frame(expand.grid(re=re, fe=fe.me))#create data frame with one column for each combination of fixed main and random effect
	to.do$re=as.character(to.do$re)#reformat to character (for later addressing by name)
	to.do$fe=as.character(to.do$fe)#reformat to character (for later addressing by name)
	res.detailed=lapply(1:nrow(to.do), function(xrow){#create detailed results by lapply-ing over the rows of to.do
			table(data[,to.do$re[xrow]], data[,to.do$fe[xrow]])#cross tabulate the respective fixed and random effect
	})
	names(res.detailed)=paste(to.do$fe, to.do$re, sep="_within_")#name it
	res.summary=lapply(1:nrow(to.do), function(xrow){#begin with creating summary results by lapply-ing over the rows of to.do
		if(modes[to.do[xrow, "fe"]]!="factor" & !treat.covs.as.factors){#if fixed effect is not a factor
			ires=table(apply(res.detailed[[xrow]]>0, 1, sum))#determine number of levels of the random effect per number of unique cases of the fixed effect
		}else{#if fixed effect is a factor
			ires=table(apply(res.detailed[[xrow]]>1, 1, sum))#determine number of levels of the random effect per number of unique cases of the fixed effect with at least two observations
		}
		ires=c(ires, tot=nrow(res.detailed[[xrow]]))
	})
	names(res.summary)=paste(to.do$fe, to.do$re, sep="_within_")#name it
	#append 'factor' or 'covariate' to the names:
	names(res.summary)[modes[to.do$fe]=="factor"]=paste(names(res.summary)[modes[to.do$fe]=="factor"], "(factor)", sep=" ")
	names(res.summary)[!modes[to.do$fe]=="factor"]=paste(names(res.summary)[!modes[to.do$fe]=="factor"], "(covariate)", sep=" ")
	to.do=data.frame(expand.grid(re=re, int=setdiff(model.terms, fe.me)))#create data frame with one row for each combination of interaction and random effect
	if(nrow(to.do)>0){
		to.do[, "int"]=as.character(to.do[, "int"])#reformat to character (for later addressing by name)
		to.do[, "re"]=as.character(to.do[, "re"])#reformat to character (for later addressing by name)
		to.add=lapply(1:nrow(to.do), function(xrow){#treat combinations of fixed and random effect by lapply-ing over the rows of to.do
			iterms=unlist(strsplit(to.do[xrow, "int"], split=":", fixed=T))#determine main effects involved in the interaction...
			imodes=modes[iterms]#... and their classes
			all.tabs=lapply(1:length(iterms), function(yrow){#for each term
				#if its not a factor, determine determine the number of unique cases of the fixed effect per level of the random effect
				#otherwise, determine per level of thee random effect the number of levels of the fixed effect with at least two observations
				apply(res.detailed[[paste(c(iterms[yrow], to.do[xrow, "re"]), collapse="_within_")]]>ifelse(imodes[iterms[yrow]]=="factor" | treat.covs.as.factors, 1, 0), 1, sum)
			})
			all.tabs=matrix(unlist(all.tabs), ncol=length(all.tabs), byrow=F)#reformat to matrix (one row per level of the random effect, one column per term)
			all.tabs=aggregate(1:nrow(all.tabs), c(data.frame(all.tabs)), length)#summarize it ((wrt the number of levels of the random effec per combination of values)
			colnames(all.tabs)=c(iterms, paste(c("n", to.do[xrow, "re"]), collapse="."))#name it
			return(all.tabs)
		})
		names(to.add)=gsub(x=paste(to.do$int, to.do$re, sep="_within_"), pattern=":", replacement="_", fixed=T)#name it
		res.summary=c(res.summary, to.add)#and append to.add to res.summary
	}
	#add columns with the dummy coded factor levels to data:
	to.code=fe.me[modes[fe.me]=="factor"]#determine factors among the fixed effects:
	if(length(to.code)>0){
		coded=lapply(to.code, function(xc){#for each factor
			lapply(levels(data[, xc])[-1], function(xl){#for all levels except the reference level
				as.numeric(data[, xc]==xl)#code it
			})
		})
		coded=matrix(unlist(coded), nrow=nrow(data), byrow=F)#reformat to matrix
		xnames=unlist(lapply(to.code, function(xc){#determine column names to be given to the matrix...
			paste(xc, levels(data[, xc])[-1], sep=".")
		}))
		colnames(coded)=xnames#... and use 'm
		data=data.frame(data, coded)#and append 'm to data
	}else{
		xnames=""
	}
	#now the model expression wrt random slopes; first no correllations between random slopes and intercepts:
	pot.terms=c(xnames, fe.me[modes[fe.me]!="factor"])#create vector with names of dummy variables and names of fixed main effects not being factors
	pot.terms=outer(pot.terms, re, Vectorize(function(x, y){#create matrix with separate model term for each combination of fixed and random effect
		paste(c("(0+", x, "|", y, ")"), collapse="")
	}))
	pot.terms=paste(c(t(pot.terms)), collapse="+")#put them all in a single entry
	#now the random slopes part of the model including random intercepts and slopes and their correlation:
	pot.terms.with.corr=paste(c(xnames, fe.me[modes[fe.me]!="factor"]), collapse="+")#get all fixed effects terms together...
	pot.terms.with.corr=paste(paste("(1+", pot.terms.with.corr, "|", re, ")", sep=""), collapse="+")#... and paste random effects, brackets and all that 
		#(and everything in a single entry)
	return(list(detailed=res.detailed, summary=res.summary, data=data, pot.terms=pot.terms, pot.terms.with.corr=pot.terms.with.corr))
}
```
