## Supplementary material for "Male Guinea baboons are oblivious to their females’ whereabouts": Data Analyses: drop1_para_coxme.r

```
drop1p<-function(model.res, para=F, data=NULL, n.cores=c("all-1", "all"), to.del=NULL, return.model.results=F, load.lib=T, lib.loc=.libPaths()){
	#last changed July 6 2017 (minor fix re number cores used)
	##determine terms that can be dropped:
	xcall=as.character(model.res$call)[2]
	#xcall=gsub(x=xcall, pattern="\n", replacement=T)
	model.terms=attr(terms(as.formula(xcall)), "term.labels")
	model.terms=model.terms[!grepl(x=model.terms, pattern="|", fixed=T)]
	model.terms=strsplit(model.terms, split=":", fixed=T)
	if(length(to.del)==0){
		to.del=unlist(lapply(1:length(model.terms), function(m1){
			xx=unlist(lapply((1:length(model.terms))[-m1], function(m2){
				length(intersect(model.terms[[m1]], model.terms[[m2]]))==length(model.terms[[m1]])
			}))
			sum(xx)>0
		}))
		model.terms=model.terms[!to.del]
		model.terms=unlist(lapply(model.terms, paste, collapse=":"))
	}else{
		model.terms=to.del
	}
	#if data are handed over
	if(length(data)>0){
		#check whether all columns needed are in them
		#figure out names of the variables involved:
		model.terms2=attr(terms(as.formula(xcall)), "term.labels")
		model.terms2=model.terms2[!grepl(x=model.terms2, pattern="|", fixed=T)]
		#model.terms2=gsub(x=model.terms2, pattern="||", replacement="|", fixed=T)
		#model.terms2=gsub(x=model.terms2, pattern="|", replacement="+", fixed=T)
		model.terms2=gsub(x=model.terms2, pattern=":", replacement="+", fixed=T)
		model.terms2=gsub(x=model.terms2, pattern="*", replacement="+", fixed=T)
		model.terms2=gsub(x=model.terms2, pattern=" ", replacement="", fixed=T)
		model.terms2=gsub(x=model.terms2, pattern="I(", replacement="", fixed=T)
		model.terms2=gsub(x=model.terms2, pattern="^2)", replacement="", fixed=T)
		model.terms2=gsub(x=model.terms2, pattern="^3)", replacement="", fixed=T)
		model.terms2=unique(unlist(strsplit(model.terms2, split="+",fixed=T)))
		model.terms2=model.terms2[!model.terms2%in%c("0", "1")]
		#deal with cbind response:
		if(substr(x=as.character(xcall), start=1, stop=6)=="cbind("){
			xx=unlist(strsplit(as.character(xcall), split="~", fixed=T))[1]
			xx=gsub(xx, pattern="cbind(", replacement="", fixed=T)
			xx=gsub(xx, pattern=")", replacement="", fixed=T)
			xx=gsub(xx, pattern=" ", replacement="", fixed=T)
			model.terms2=c(model.terms2, unlist(strsplit(xx, split=",", fixed=T)))
		}
		#deal with sine/cosine included into model:
		model.terms2=model.terms2[!substr(model.terms2, start=1, stop=4)=="sin("]
		model.terms2=model.terms2[!substr(model.terms2, start=1, stop=4)=="cos("]
		
		#still missing: offset term and weights, squared terms...
		if(any(!model.terms2%in%names(data))){
			xx=model.terms2[!model.terms2%in%names(data)]
			stop(paste(c(paste(xx, collapse=", "), ifelse(length(xx)==1, "is ", "are "), "missing in data"), collape=""))
		}
	}
	model.terms2=model.terms[!grepl(x=model.terms, pattern="|", fixed=T)]
	##prepare data:
	ii.data=data
	#all.models=c(xcall, paste(xcall, model.terms, sep="-"))
	all.models=paste(xcall, model.terms, sep="-")
	##define function doing the model comparison:
	model.fun<-function(i.model, idata){
		i.model=as.formula(i.model)
		i.res=coxme(i.model, data=idata)
		return(i.res)
	}
	##run all models:
  if(para){
    require(parallel)
    n.cores=n.cores[1]
    cl <- makeCluster(getOption("cl.cores", detectCores()))
		if(n.cores=="all"){
			n.cores=length(cl)
		}else if(n.cores=="all-1"){
			if(n.cores=="all-1"){n.cores=length(cl)-1}#else{n.cores=length(cl)}
		}
		n.cores=min(c(n.cores, length(all.models)))
		cl=cl[1:n.cores]
    parLapply(cl=cl, 1:length(cl), fun=function(x){
      return(invisible(library(coxme, lib.loc=lib.loc)))
    })
    x.all.res=parLapply(cl=cl, X=all.models, fun=model.fun, idata=ii.data)
    parLapply(cl=cl, X=1:length(cl), fun=function(x){invisible(rm(list=ls()))})
    stopCluster(cl)
  }else{
		if(load.lib){library(coxme, lib.loc=lib.loc)}
		x.all.res=lapply(X=all.models, FUN=model.fun, idata=ii.data)
  }
	all.tests=lapply(x.all.res, function(x){
		c(x$loglik["Integrated"], length(fixef(x)))
	})
	all.tests=matrix(unlist(all.tests), nrow=length(all.tests), byrow=T)
	colnames(all.tests)=c("loglik", "df")
	all.tests=cbind(all.tests, chi=-2*(all.tests[, "loglik"]-model.res$loglik["Integrated"]))
	all.tests=cbind(all.tests, chi.df=length(fixef(model.res))-all.tests[, "df"])
	all.tests=cbind(all.tests, P=pchisq(q=all.tests[, "chi"], df=all.tests[, "chi.df"], lower.tail=F))
	rownames(all.tests)=model.terms
  return(all.tests)
}
```
