## Supplementary material for "Male Guinea baboons are oblivious to their females’ whereabouts": Data Analyses: ExperimentsAnalyses.R

```
###############################################################
# Male Guinea baboon tracking of female whereabouts	        #
# 	     								        #
# Dominique Treschnak,Dietmar Zinner, Julia Fischer	        #	
#									        #
# Script for statitical analyses and plotting of main results #
###############################################################

# Set working direction to the folder containing data files and all scripts
# Necessary packages: lme4, survival, coxme, lmerTest
# setwd()

# Load needed packages and source necessary functions
library(lme4)
source("diagnostic_fcns.r")
source("glmm_stability.r")
source("boot_glmm.r")

##### Experiment 1(individual recognition) #####

### Load dataset: Experiment 1 
dat<-read.csv("Summary_Looking_B.csv",stringsAsFactors=T)

# Load needed packages and source necessary functions
library(lme4)
source("diagnostic_fcns.r")
source("glmm_stability.r")
source("boot_glmm.r")

## Gaussian model for Looking Duration

# Log-transformation of Looking Duration
dat$log.First_Look_Dur<-log(dat$First_Look_Dur+1)

# Calculate Gaussian model
gauss.dur.m<-lmer(log.First_Look_Dur~Condition+Nr_Females+(1|MaleID)+(1|CallID)+(1|PartnerID),data=dat,REML=F)

# Check of model assumptions using diagnostics.plot() function
diagnostics.plot(gauss.dur.m)

# Check model stability using glmm.model.stab() function (recalculates model estimates removing individual level of random effects one at a time)
gauss.dur.m.stab <- glmm.model.stab(model.res = gauss.dur.m)
# Inspect minimum and maximum model estimates (as table and in plot)
gauss.dur.m.stab$summary[,-1]
m.stab.plot(gauss.dur.m.stab$summary[,-1])

# Parametric bootstrapping of confidence interval for model estimates
gauss.dur.m.boot.full<-boot.glmm.pred(model.res=gauss.dur.m,use=dat$Condition)
round(gauss.dur.m.boot.full$ci.estimates,3)
m.stab.plot(gauss.dur.m.boot.full$ci.estimates)5

# Calculating null model without main effect: Condition
gauss.dur.m0<-lmer(log.First_Look_Dur~Nr_Females+(1|MaleID)+(1|CallID)+(1|PartnerID),data=dat,REML=F)

# Full model comparison using anova() function
as.data.frame(anova(gauss.dur.m0,gauss.dur.m,test="Chisq"))

# Calculating p-values of individual fixed effects using drop1() function
as.data.frame(drop1(gauss.dur.m,test="Chisq"))

# Model summary
summary(gauss.dur.m)

# Calculate t and p values using lmer() function of lmerTest package
library(lmerTest)
gauss.dur.m.reml=lmer(log.First_Look_Dur~Condition+Nr_Females+(1|MaleID)+(1|CallID)+(1|PartnerID), data=dat, REML=T)
round(summary(gauss.dur.m.reml)$coeff,3)
detach("package:lmerTest")

### Create Plot for Looking Duration of Experiment 2 (individual recognition)

# It is necessary to run the transformations of the outcome variables (from original model) before plotting!

# Create new standardised variable for Number of Females, allows to plot the effect of the main
# effect Condition for a male with an average Number of Females
st.Nr_Females<-as.vector(scale(dat$Nr_Females))

# Calculate new model with standardised Nr_Females
plot.gauss.dur.m<-lmer(log.First_Look_Dur~Condition+st.Nr_Females+(1|MaleID)+(1|CallID)+(1|PartnerID),data=dat,REML=F)

# Parametric bootstrapping of confidence interval for model estimates with standardised Nr_Females
plot.gauss.dur.m.boot.full<-boot.glmm.pred(model.res=plot.gauss.dur.m,use=dat$Condition)
round(plot.gauss.dur.m.boot.full$ci.estimates,3)

# Plotting:

# Create objects for model prediction and bootstrapped values for confidence interval
dur.lowCI_nuf<-(plot.gauss.dur.m.boot.full$ci.predict[1,4])
dur.upCI_nuf<-(plot.gauss.dur.m.boot.full$ci.predict[1,5])
dur.est_nuf<-(plot.gauss.dur.m.boot.full$ci.predict[1,3])
dur.lowCI_uf<-(plot.gauss.dur.m.boot.full$ci.predict[2,4])
dur.upCI_uf<-(plot.gauss.dur.m.boot.full$ci.predict[2,5])
dur.est_uf<-(plot.gauss.dur.m.boot.full$ci.predict[2,3])
# Prepare plotting window
par(mar=c(3,3,0.2,0.2),mgp=c(1.7,0.3,0),tcl=-0.15,las=1)
# Prepare axis
plot.default(x=dat$Condition,y=dat$First_Look_Dur,xaxt="n",xlim=c(0.5,2.5),xaxs="i",col=grey(level=0.4,alpha=0.4),cex=1.5,pch=19,type="n",
xlab="Conditions",ylab="First look duration [s]",cex.lab=1.4)
mtext(text=c("Non-unit-female","Unit-female"),side=1,at=1:2,line=0.2,cex=1.3)
# Calculate and plot lines connecting data points of same individuals tested in both conditions
xx<-tapply(X=dat$First_Look_Dur,INDEX=list(dat$MaleID,dat$Condition),FUN=mean)
segments(x0=1,x1=2,y0=xx[,1],y1=xx[,2])
# Plotting of individual data points
points(x=dat$Condition,y=dat$First_Look_Dur,cex=1.5,pch=19,col="white")
points(x=dat$Condition,y=dat$First_Look_Dur,col=grey(level=0.4,alpha=0.4),cex=1.5,pch=19)
# Plotting of model predictions and confidence interval
arrows(y0=exp(dur.lowCI_nuf),y1=exp(dur.upCI_nuf),x0=1,x1=1,angle=90,length=0.15,code=3,lwd=2);segments(x0=0.97,x1=1.03,y0=exp(dur.est_nuf),y1=exp(dur.est_nuf),lwd=2)
arrows(y0=exp(dur.lowCI_uf),y1=exp(dur.upCI_uf),x0=2,x1=2,angle=90,length=0.15,code=3,lwd=2);segments(x0=1.97,x1=2.03,y0=exp(dur.est_uf),y1=exp(dur.est_uf),lwd=2)


## Survival model for Looking Latency

# Load needed packages and source necessary functions
library(survival)
library(coxme)
source("coxme_stab.r")

# Create survival object using Surv() function (censors Looking Latencies >2.5 s)
dat$rv<-dat$First_Look_Lat
dat$rv=Surv(time=dat$rv,event=dat$First_Look_Lat<2.5)

# Calculate survival model
surv.lat.m<-coxme(rv~Condition+Nr_Females+(1|MaleID)+(1|CallID)+(1|PartnerID),data=dat)

# Calculating null model without main effect: Condition
surv.lat.m0<-coxme(rv~Nr_Females+(1|MaleID)+(1|CallID)+(1|PartnerID),data=dat)

# Full model comparison using anova() function
as.data.frame(anova(surv.lat.m,surv.lat.m0))

# Calculating p-values of individual fixed effects using drop1p() function
source("drop1_para_coxme.r")
drop1p(model.res=surv.lat.m, para=F, data=dat, n.cores=c("all-1", "all"),
to.del=NULL, return.model.results=F, load.lib=T, lib.loc=.libPaths())


# Check model stability using glmm.model.stab() function (recalculates model estimates removing individual level of random effects one at a time)
surv.lat.m.stab<-coxme.stab(mres=surv.lat.m,data=dat,formula=rv~Condition+Nr_Females+(1|MaleID)+(1|CallID)+(1|PartnerID))
# Inspect minimum and maximum model estimates (as table and in plot)
surv.lat.m.stab$summary[,-1]
m.stab.plot(surv.lat.m.stab$summary)

# Model summary
summary(surv.lat.m)


##### Experiment 2 (spatial monitoring) #####

### Load dataset: Experiment 2 
dat<-read.csv("Summary_Looking_A.csv",stringsAsFactors=T)

## Gamma model for Looking Duration

# Prepare response variable (First_Look_Dur) for use in gamma model; values of 0 become 0.01
dat$tr.First_Look_Dur<-dat$First_Look_Dur
dat$tr.First_Look_Dur[dat$tr.First_Look_Dur==0]<-0.01

# Calculate Gamma model
gamma.dur.m<-glmer(tr.First_Look_Dur~Condition+Nr_Females+(1|MaleID),data=dat,family=Gamma(link="log"))

# Check for overdispersion using overdisp.test() function
overdisp.test(gamma.dur.m)

# Check model stability using glmm.model.stab() function (recalculates model estimates removing individual level of random effects one at a time)
gamma.dur.m.stab <- glmm.model.stab(model.res = gamma.dur.m)
# Inspect minimum and maximum model estimates (as table and in plot)
gamma.dur.m.stab$summary[,-1]
m.stab.plot(gamma.dur.m.stab$summary[,-1])

# Parametric bootstrapping of confidence interval for model estimates
gamma.dur.m.boot.full<-boot.glmm.pred(model.res=gamma.dur.m,use=dat$Condition)
round(gamma.dur.m.boot.full$ci.estimates,3)

# Calculating null model without main effect: Condition
gamma.dur.m0<-glmer(tr.First_Look_Dur~Nr_Females+(1|MaleID),data=dat,family=Gamma(link="log"))

# Full model comparison using anova() function
as.data.frame(anova(gamma.dur.m0,gamma.dur.m,test="Chisq"))

# Calculating p-values of individual fixed effects using drop1() function
as.data.frame(drop1(gamma.dur.m,test="Chisq"))

# Model summary
summary(gamma.dur.m)$coeff
summary(gamma.dur.m)$varcor

### Create Plot for Looking Duration of Experiment 1 (spatial monitoring)

# It is necessary to run the transformations of the outcome variables (from original model) before plotting!

# Create new standardised variable for Number of Females, allows to plot the effect of the main
# effect Condition for a male with an average Number of Females

# Standardising variable Nr_Females
st.Nr_Females<-as.vector(scale(dat$Nr_Females))

# Calculate new model with standardised Nr_Females
plot.gamma.dur.m<-glmer(tr.First_Look_Dur~Condition+st.Nr_Females+(1|MaleID),data=dat,family=Gamma(link="log"))

# Parametric bootstrapping of confidence interval for model estimates with standardised Nr_Females
plot.gamma.dur.m.boot.full<-boot.glmm.pred(model.res=plot.gamma.dur.m,use=dat$Condition)
round(plot.gamma.dur.m.boot.full$ci.estimates,3)

# Plotting:

# Create objects for model prediction and bootstrapped values for confidence interval 
dur.lowCI_con<-(plot.gamma.dur.m.boot.full$ci.predict[1,4])
dur.upCI_con<-(plot.gamma.dur.m.boot.full$ci.predict[1,5])
dur.est_con<-(plot.gamma.dur.m.boot.full$ci.predict[1,3])
dur.lowCI_inc<-(plot.gamma.dur.m.boot.full$ci.predict[2,4])
dur.upCI_inc<-(plot.gamma.dur.m.boot.full$ci.predict[2,5])
dur.est_inc<-(plot.gamma.dur.m.boot.full$ci.predict[2,3])
# Prepare plotting window
par(mar=c(3,3,0.2,0.2),mgp=c(1.7,0.3,0),tcl=-0.15,las=1)
# Prepare axis
plot.default(x=dat$Condition,y=dat$First_Look_Dur,xaxt="n",xlim=c(0.5,2.5),xaxs="i",col=grey(level=0.4,alpha=0.4),cex=1.5,pch=19,type="n",
xlab="Conditions",ylab="First look duration [s]",cex.lab=1.4)
mtext(text=c("Consistent","Inconsistent"),side=1,at=1:2,line=0.2,cex=1.3)
# Calculate and plot lines connecting data points of same individuals tested in both conditions
xx<-tapply(X=dat$First_Look_Dur,INDEX=list(dat$RunNr,dat$Condition),FUN=mean)
segments(x0=1,x1=2,y0=xx[,1],y1=xx[,2])
# Plotting of individual data points
points(x=dat$Condition,y=dat$First_Look_Dur,cex=1.5,pch=19,col="white")
points(x=dat$Condition,y=dat$First_Look_Dur,col=grey(level=0.4,alpha=0.4),cex=1.5,pch=19)
# Plotting of model predictions and confidence interval
arrows(y0=dur.lowCI_con,y1=dur.upCI_con,x0=1,x1=1,angle=90,length=0.15,code=3,lwd=2);segments(x0=0.97,x1=1.03,y0=dur.est_con,y1=dur.est_con,lwd=2)
arrows(y0=dur.lowCI_inc,y1=dur.upCI_inc,x0=2,x1=2,angle=90,length=0.15,code=3,lwd=2);segments(x0=1.97,x1=2.03,y0=dur.est_inc,y1=dur.est_inc,lwd=2)

## Survival model for Looking Latency

# Load needed packages and source necessary functions (please install necessary packages)
library(survival)
library(coxme)
source("coxme_stab.r")

# Create survival object using Surv() function (censors Looking Latencies >2.5 s) 
dat$rv<-dat$First_Look_Lat
dat$rv=Surv(time=dat$rv,event=dat$First_Look_Lat<2.5)

# Calculate survival model
surv.lat.m<-coxme(rv~Condition+Nr_Females+(1|MaleID),data=dat)

# Calculating null model without main effect: Condition
surv.lat.m0<-coxme(rv~Nr_Females+(1|MaleID),data=dat)

# Full model comparison using anova() function
as.data.frame(anova(surv.lat.m0,surv.lat.m))

# Calculating p-values of individual fixed effects using drop1p() function
source("drop1_para_coxme.r")
drop1p(model.res=surv.lat.m, para=F, data=dat, n.cores=c("all-1", "all"),
to.del=NULL, return.model.results=F, load.lib=T, lib.loc=.libPaths())

# Check model stability using coxme.stab() function (recalculates model estimates removing individual level of random effects one at a time)
surv.lat.m.stab<-coxme.stab(mres=surv.lat.m,data=dat,formula=rv~Condition+Nr_Females+(1|MaleID))
# Inspect minimum and maximum model estimates (as table and in plot)
surv.lat.m.stab$summary[,-1]
m.stab.plot(surv.lat.m.stab$summary)

# Non-parametric bootstrapping of confidence interval for model estimates

nboot=1000
toran<-lapply(X=levels(dat$MaleID),FUN=function(x){
	which(dat$MaleID==x)
})
allest<-matrix(NA,nrow=nboot,ncol=length(coef(surv.lat.m)))
colnames(allest)<-names(coef(surv.lat.m))
for (i in 1:nboot){
	xboot<-sample(1:length(toran),size=length(toran),replace=T)
	bootdata=dat[unlist(toran[xboot]),]
	bootdata$MaleID=as.factor(rep(x=1:length(xboot),unlist(lapply(X=toran[xboot],FUN=length))))
	mb<-try(coxme(rv~Condition+Nr_Females+(1|MaleID),data=bootdata),silent=T)
	if (class(mb)[[1]]!="try-error"){
		allest[i,]=coef(mb)
	}
}	 

res.lat<-apply(X=allest,MARGIN=2,FUN=quantile,probs=c(0.025,0.975),na.rm=T)
sum(!is.na(allest[,1]))
res.lat

# Model summary
summary(surv.lat.m)

## Gauss model for Vigilance time

# Square-transformation of Vigilance time
dat$sq.Vigilance<-sqrt(dat$Vigilance)

# Calculate Gaussian model
v.m<-lmer(sq.Vigilance~Condition+Nr_Females+(1|MaleID),data=dat,REML=F)

# Check of model assumptions using diagnostics.plot() function
diagnostics.plot(v.m)

# Check model stability using glmm.model.stab() function (recalculates model estimates removing individual level of random effects one at a time)
v.m.stab <- glmm.model.stab(model.res = v.m)
# Inspect minimum and maximum model estimates (as table and in plot)
v.m.stab$summary[,-1]
m.stab.plot(v.m.stab$summary[,-1])

# Parametric bootstrapping of confidence interval for model estimates
v.m.boot.full<-boot.glmm.pred(model.res=v.m,use=dat$Condition)
round(v.m.boot.full$ci.estimates,3)

# Calculating null model without main effect: Condition
v.m0<-lmer(sq.Vigilance~Nr_Females+(1|MaleID),data=dat,REML=F)

# Full model comparison using anova() function
as.data.frame(anova(v.m0,v.m,test="Chisq"))

# Calculating p-values of individual fixed effects using drop1() function
as.data.frame(drop1(v.m,test="Chisq"))

# Model summary
summary(v.m)

# Calculate t and p values using lmer() function of lmerTest package; please install necessary packages
library(lmerTest)
v.m.reml=lmer(sq.Vigilance~Condition+Nr_Females+(1|MaleID), data=dat, REML=T)
round(summary(v.m.reml)$coeff,3)
detach("package:lmerTest")

##### Experiment 2 (spatial monitoring) - Alternative models, including proximity of other baboon parties (Mingled: Y/N) #####

## Gamma model for Looking Duration (Alternative)

# Prepare response variable (First_Look_Dur) for use in gamma model; values of 0 become 0.01
dat$tr.First_Look_Dur<-dat$First_Look_Dur
dat$tr.First_Look_Dur[dat$tr.First_Look_Dur==0]<-0.01

# Calculate Gamma model
gamma.dur.alt.m<-glmer(tr.First_Look_Dur~Condition+Nr_Females+Mingled+(1|MaleID),data=dat,family=Gamma(link="log"))

# Check for overdispersion using overdisp.test() function
overdisp.test(gamma.dur.alt.m)

# Check model stability using glmm.model.stab() function (recalculates model estimates removing individual level of random effects one at a time)
gamma.dur.alt.m.stab <- glmm.model.stab(model.res = gamma.dur.alt.m)
# Inspect minimum and maximum model estimates (as table and in plot)
gamma.dur.alt.m.stab$summary[,-1]
m.stab.plot(gamma.dur.alt.m.stab$summary[,-1])

# Parametric bootstrapping of confidence interval for model estimates
gamma.dur.alt.m.boot.full<-boot.glmm.pred(model.res=gamma.dur.alt.m,use=dat$Condition)
round(gamma.dur.alt.m.boot.full$ci.estimates,3)
m.stab.plot(gamma.dur.alt.m.boot.full$ci.estimates)

# Calculating null model without main effect: Condition
gamma.dur.alt.m0<-glmer(tr.First_Look_Dur~Nr_Females+Mingled+(1|MaleID),data=dat,family=Gamma(link="log"))

# Full model comparison using anova() function
as.data.frame(anova(gamma.dur.alt.m0,gamma.dur.m,test="Chisq"))

# Calculating p-values of individual fixed effects using drop1() function
as.data.frame(drop1(gamma.dur.alt.m,test="Chisq"))

# Model summary
summary(gamma.dur.alt.m)

## Survival model for Looking Latency (Alternative)

# Load needed packages and source necessary functions
library(survival)
library(coxme)
source("coxme_stab.r")

# Create survival object using Surv() function (censors Looking Latencies >2.5 s)
dat$rv<-dat$First_Look_Lat
dat$rv=Surv(time=dat$rv,event=dat$First_Look_Lat<2.5)

# Calculate survival model
surv.lat.alt.m<-coxme(rv~Condition+Nr_Females+Mingled+(1|MaleID),data=dat)

# Calculating null model without main effect: Condition
surv.lat.alt.m0<-coxme(rv~Nr_Females+Mingled+(1|MaleID),data=dat)

# Full model comparison using anova() function
as.data.frame(anova(surv.lat.alt.m0,surv.lat.alt.m))

# Calculating p-values of individual fixed effects using drop1p() function
source("drop1_para_coxme.r")
tests<-drop1p(model.res=surv.lat.alt.m, para=F, data=dat, n.cores=c("all-1", "all"),
to.del=NULL, return.model.results=F, load.lib=T, lib.loc=.libPaths())
tests

# Check model stability using glmm.model.stab() function (recalculates model estimates removing individual level of random effects one at a time)
surv.lat.alt.m.stab<-coxme.stab(mres=surv.lat.alt.m,data=dat,formula=rv~Condition+Nr_Females+Mingled+(1|MaleID))
# Inspect minimum and maximum model estimates (as table and in plot)
surv.lat.alt.m.stab$summary[,-1]
m.stab.plot(surv.lat.alt.m.stab$summary)

# Non-parametric bootstrapping of confidence interval for model estimates
nboot=1000
toran<-lapply(X=levels(dat$MaleID),FUN=function(x){
	which(dat$MaleID==x)
})

allest<-matrix(NA,nrow=nboot,ncol=length(coef(surv.lat.alt.m)))
colnames(allest)<-names(coef(surv.lat.alt.m))
for (i in 1:nboot){
	xboot<-sample(1:length(toran),size=length(toran),replace=T)
	bootdata=dat[unlist(toran[xboot]),]
	bootdata$MaleID=as.factor(rep(x=1:length(xboot),unlist(lapply(X=toran[xboot],FUN=length))))
	mb<-try(coxme(rv~Condition+Nr_Females+Mingled+(1|MaleID),data=bootdata),silent=T)
	if (class(mb)[[1]]!="try-error"){
		allest[i,]=coef(mb)
	}
}	 

res.lat.alt<-apply(X=allest,MARGIN=2,FUN=quantile,probs=c(0.025,0.975),na.rm=T)
res.lat.alt

# Model summary
summary(surv.lat.alt.m)

## Gaussian model for Vigilance time (Alternative)

# Square-transformation of Vigilance time
dat$sq.Dur<-sqrt(dat$Vigilance)

# Calculate Gaussian model
v.alt.m<-lmer(sq.Dur~Condition+Nr_Females+Mingled+(1|MaleID),data=dat,REML=F)

# Check of model assumptions using diagnostics.plot() function
diagnostics.plot(v.alt.m)

# Check model stability using glmm.model.stab() function (recalculates model estimates removing individual level of random effects one at a time)
v.alt.m.stab <- glmm.model.stab(model.res = v.alt.m)
# Inspect minimum and maximum model estimates (as table and in plot)
v.alt.m.stab$summary[,-1]
m.stab.plot(v.alt.m.stab$summary[,-1])

# Parametric bootstrapping of confidence interval for model estimates
v.alt.m.boot.full<-boot.glmm.pred(model.res=v.alt.m,use=dat$Condition)
round(v.alt.m.boot.full$ci.estimates,3)
m.stab.plot(v.alt.m.boot.full$ci.estimates)

# Calculating null model without main effect: Condition
v.alt.m0<-lmer(sq.Dur~Nr_Females+Mingled+(1|MaleID),data=dat,REML=F)

# Full model comparison using anova() function
as.data.frame(anova(v.alt.m0,v.alt.m,test="Chisq"))

# Calculating p-values of individual fixed effects using drop1() function
as.data.frame(drop1(v.alt.m,test="Chisq"))

# Model summary
summary(v.alt.m)

# Calculate t and p values using lmer() function of lmerTest package
library(lmerTest)
v.alt.m.reml=lmer(sq.Dur~Condition+Nr_Females+Mingled+(1|MaleID), data=dat, REML=T)
round(summary(v.alt.m.reml)$coeff,3)
detach("package:lmerTest")
```
