## Supplementary material for "Male Guinea baboons are oblivious to their females’ whereabouts": Data Analyses: glmm_stability.r

```
glmm.model.stab<-function(model.res, contr=NULL, ind.cases=F, para=F, data=NULL, use=NULL, n.cores=c("all-1", "all"), save.path=NULL, load.lib=T, lib.loc=.libPaths()){
	print("please carefully evaluate whether the result makes sense, and if not, please contact me")
	#function determining stability of GLMMs (run using lmer or glmer) by excluding levels of random effects, one at a time;
	#supports
		#weights, offset terms and random slopes;
	#does not support
		#correlations between random intercepts and random slopes
		#any terms calculated in the model formula (e.g., log, function I, etc.); but interactions do work
	#latest additions/modifications:
		#copes also with data sets where a level of a (fixed effects) factor is entirely dropped from the data
		#new way of catching warnings (are contained in the detailed output table)
		#includes sample size in the detailed output table
		#catches errors
		#use: an argument taking the names of the random effects for which model stability should be evaluated
			#(useful for models with a random effect having a unique case for each level of the data)
	#written by Roger Mundry
	#modified April 2016 (added dealing with glmer.nb; fixed a bug in the output of the random effects of tthe original model)
	#last modified Mar 2017 (added dealing with fixed effects factor levels dropped when dropping levels of random effects)
	#last modified June 14 2017 (fixed dealing with random effects comprising correlation parameters)
	#last modified July 06 2017 (fixed small bug happening when model revealed an error)
	if(load.lib){library(lme4, lib.loc=lib.loc)}
	n.cores=n.cores[1]
  model.eq=as.formula(as.character(model.res@call)[2])
	if(class(model.res)[1]=="lmerMod"){
    REML=model.res@resp$REML==1
    xfam="gaussian"
  }else{
		xfam=model.res@resp$family$family
		if(model.res@resp$family$family=="Gamma"){
			xfam=model.res@resp$family
		}
  }
  if(grepl(x=xfam[[1]], pattern="Negative Binomial", fixed=T)){
		xfam="neg.bin"
	}
  weights=model.res@resp$weights
  if(length(data)==0){
		ii.data=model.res@frame
		offs.col=grep(x=names(ii.data), pattern="offset(", fixed=T)
		if(length(offs.col)>0){
			for(i in offs.col){
				#ii.data[,i]=exp(ii.data[,i])
				names(ii.data)[i]=substr(x=names(ii.data)[i], start=8, stop=nchar(names(ii.data)[i]) -1)
			}
		}
		wght.col=grep(x=names(ii.data), pattern="(weights)", fixed=T)
		if(length(wght.col)>0){
			names(ii.data)[wght.col]=gsub(x=names(ii.data)[wght.col], pattern="(", replacement="", fixed=T)
			names(ii.data)[wght.col]=gsub(x=names(ii.data)[wght.col], pattern=")", replacement="", fixed=T)
		}else{
			ii.data$weights=weights
		}
	}else{
		ii.data=data.frame(data, weights)
	}
	if(substr(as.character(model.eq)[2], start=1, stop=6)=="cbind("){
		ii.data$weights=1
	}
  #if(xfam=="binomial"){
  #   response=as.character(model.eq)[2]
  #   if(sum(names(ii.data)==response)==0)
  #}
  ranefs=names(ranef(model.res))
	if(length(use)==0){use=ranefs}
	ranefs=ranefs[ranefs%in%use]
  xlevels=lapply(ranefs, function(x){return(as.vector(unique(ii.data[ ,x])))})
  ranefs=rep(ranefs, unlist(lapply(xlevels, length)))
  to.do=cbind(ranefs, unlist(xlevels))
  if(ind.cases){
    ii.data=data.frame(ii.data, ic=as.factor(1:nrow(ii.data)))
    to.do=rbind(to.do, cbind("ic", levels(ii.data$ic)))
  }
	keepWarnings <- function(expr) {
		localWarnings <- list()
		value <- withCallingHandlers(expr,
			warning = function(w) {
				localWarnings[[length(localWarnings)+1]] <<- w
				invokeRestart("muffleWarning")
			}
		)
		list(value=value, warnings=localWarnings)
	}
	get.ranef<-function(x){#function returning sd associated with random effect in a nicely named vector
		x=as.data.frame(x)
		y=lapply(strsplit(as.character(x$grp), split=""), function(y){
			yy=unlist(lapply(strsplit(names(ranef(model.res)), split=""), function(z){
				if(length(z)<=length(y)){
					y=y[1:length(z)]
					return(paste(y[y==z], collapse=""))
				}else{
					return("")
				}
			}))
			yy=intersect(yy, names(ranef(model.res)))
			if(length(yy)==0){yy=paste(y, collapse="")}
			return(yy)
		})
		x$grp=y
		ires=x$sdcor
		#x$var2[is.na(x$var2)]=""
		names(ires)=apply(x[, 1:3], 1, paste, collapse="@")
		names(ires)[x$grp=="Residual"]="Residual"
		return(ires)
	}
	if(length(contr)==0){
		if(class(model.res)[1]=="lmerMod"){
			contr=lmerControl()
		}else{	
			contr=glmerControl()
		}
	}
  ifun=function(x, model.res, to.do, ii.data, contr, get.ranef){
    #sel.ii.data=subset(ii.data, ii.data[,to.do[x, 1]]!=to.do[x, 2])
    sel.ii.data=ii.data[ii.data[,to.do[x, 1]]!=to.do[x, 2], ]
		if(class(model.res)[1]=="lmerMod"){
      sel.ii.res=try(lmer(model.eq, data=sel.ii.data, weights=weights, REML=REML, control=contr), silent=T)
    }else if(xfam[[1]]!="neg.bin"){#(xfam=="binomial" | xfam=="poisson"){
      sel.ii.res=try(glmer(model.eq, data=sel.ii.data, family=xfam, control=contr), silent=T)
    }else{
			sel.ii.res=try(glmer.nb(model.eq, data=sel.ii.data, control=contr), silent=T)
		}
		if(length(save.path)>0){
			est.fixed.effects=fixef(sel.ii.res)
			est.random.effects=as.data.frame(summary(sel.ii.res)$varcor)
			model.warnings=sel.ii.res$warnings
			n=length(residuals(sel.ii.res))
			what=to.do[x, ]
			save(file=paste(c(paste(c(paste(c(save.path, "m"), collapse="/"), x), collapse="_"), ".RData"), collapse=""), list=c("what", "est.fixed.effects", "est.random.effects", "model.warnings", "n"))
		}
    if(class(sel.ii.res)!="try-error"){
			#xx=try(list(fere=c(fixef(sel.ii.res$value), get.ranef(summary(sel.ii.res$value)$varcor)), N=length(residuals(sel.ii.res$value)), warnings=paste(unlist(sel.ii.res$warnings)$message, collapse="/")), silent=T)
			#if(class(xx)=="try-error"){browser()}
      return(list(fere=c(fixef(sel.ii.res), get.ranef(summary(sel.ii.res)$varcor)), N=nrow(sel.ii.res@frame), 
				warnings=
					list(
						lme4=summary(sel.ii.res)$optinfo$conv$lme4$messages,
						opt=summary(sel.ii.res)$optinfo$warnings,
						conv.code=summary(sel.ii.res)$optinfo$conv$opt
					)))
    }else{
      return(list(fere=rep(NA, length(fixef(model.res))+nrow(as.data.frame(summary(model.res)$varcor))), N=nrow(sel.ii.data), 
				warnings=list(lme4=NA, opt=NA, conv.code=NA)))
    }
  }
  if(para){
		#on.exit(expr = parLapply(cl=cl, X=1:length(cl), fun=function(x){rm(list=ls())}), add = FALSE)
		#on.exit(expr = stopCluster(cl), add = T)
    require(parallel)
    cl <- makeCluster(getOption("cl.cores", detectCores()))
		if(n.cores!="all"){
			if(n.cores=="all-1"){n.cores=length(cl)-1}
			if(n.cores<length(cl)){
				cl=cl[1:n.cores]
			}
		}
    parLapply(cl=cl, 1:length(cl), fun=function(x){
      library(lme4, lib.loc=lib.loc)
      return(invisible(""))
    })
    all.coeffs=parLapply(cl=cl, X=1:nrow(to.do), fun=ifun, model.res=model.res, to.do=to.do, ii.data=ii.data, contr=contr, get.ranef=get.ranef)
    parLapply(cl=cl, 1:length(cl), fun=function(x){
      return(rm(list=ls()))
    })
    stopCluster(cl=cl)
  }else{
		#browser()
    all.coeffs=lapply(1:nrow(to.do), ifun, model.res=model.res, to.do=to.do, ii.data=ii.data, contr=contr, get.ranef=get.ranef)
  }
  all.n=unlist(lapply(all.coeffs, function(x){x$N}))
  all.warnings=lapply(all.coeffs, function(x){
		x=sapply(x$warnings, paste, collapse="")
	})
  all.warnings=as.data.frame(matrix(unlist(all.warnings), ncol=3, byrow=T))
  colnames(all.warnings)=c("lme4.w", "opt.w", "conv.code")
	############################################################################################################################
	############################################################################################################################
	############################################################################################################################
	############################################################################################################################
	xnames=unique(unlist(lapply(all.coeffs, function(x){names(x$fere)})))
	all.coeff.mat=matrix(NA, ncol=length(xnames), nrow=length(all.coeffs))
	colnames(all.coeff.mat)=xnames
	for(i in 1:length(all.coeffs)){
		all.coeff.mat[i, names(all.coeffs[[i]]$fere)]=all.coeffs[[i]]$fere
	}
	#extract results for original model:
	orig=c(fixef(model.res), get.ranef(summary(model.res)$varcor))
	xsum=apply(all.coeff.mat, 2, range, na.rm=T)
  xsum=data.frame(what=colnames(all.coeff.mat), orig=orig[colnames(all.coeff.mat)], t(xsum))
	rownames(xsum)=as.character(xsum$what)
  colnames(to.do)=c("ranef", "level")
  xx=apply(is.na(all.coeff.mat), 1, sum)
  if(sum(xx>0 & xx<ncol(all.coeff.mat))>0){
		warning(paste(c("for", sum(xx>0 & xx<ncol(all.coeff.mat)), "subset(s) the full model could not be fitted because of fixed effects factor levels dropped from the data"), collapse=" "))
	}
	colnames(all.warnings)[1:2]=c("lme4.warnings", "opt.warnings")
  all.coeff.mat=data.frame(to.do, N=all.n, all.coeff.mat, all.warnings)
  names(xsum)[3:4]=c("min", "max")
  return(list(detailed=all.coeff.mat, summary=xsum))
}
```
